# MiTo: tracing the phenotypic evolution of somatic cell lineages via mitochondrial single-cell multi-omics

**DOI:** 10.1101/2025.06.17.660165

**Authors:** Andrea Cossa, Alberto Dalmasso, Guido Campani, Elisa Bugani, Chiara Caprioli, Noemi Bulla, Andrea Tirelli, Yinxiu Zhan, Pier Giuseppe Pelicci

**Affiliations:** Department of Experimental Oncology, European Institute of Oncology, via Adamello 16, 20139 Milan, Italy; Dipartimento di Oncologia ed Emato-Oncologia, Universita’ degli Studi di Milano, via Santa Sofia 9, 20142 Milan, Italy

## Abstract

Mitochondrial single-cell lineage tracing (MT-scLT) has recently emerged as a scalable and non-invasive tool to trace somatic cell lineages. However, the reliability and resolution of MT-scLT remains highly debated. Here, we present `MiTo`, the first end-to-end framework for robust MT-scLT data analysis. Thanks to highly-optimized algorithms and user-friendly interfaces, this modular toolkit offers unprecedented control across the entire MT-scLT workflow. Benchmarked against novel real-world datasets (375–2,757 cells; 8–216 lentiviral clones), `MiTo` outperformed state-of-the-art methods and baselines in MT-scLT data pre-processing and clonal inference. Applied to a time-resolved dataset of breast cancer evolution (>2,500 cells), `MiTo` accurately inferred ground-truth cell lineages (ARI=0.94) and cell state transitions, detected clonal fitness markers, and quantified heritability of gene regulatory networks. Comparing alternative lineage markers, `MiTo` quantified the resolution limit of existing MT-scLT systems, which currently enable reliable inference of coarse-grained cellular ancestries, but not high-resolution phylogenetic inference. In conclusion, this work provides robust tools and practical guidelines to dissect somatic evolution with single-cell multi-omics.

## Introduction

Somatic evolution, the continuous accumulation and selection of inheritable (epi)genetic mutations across an individual’s lifetime, underpins both normal tissue development and pathological processes such as cancer^1–3^. From early embryogenesis onward, mitotic divisions are accompanied by tightly regulated molecular changes and cell fate decisions, giving rise to diverse cell lineages with specialized function and heterogeneous fitness. When these developmental programs are disrupted, the same evolutionary forces can drive malignant transformation, promoting the uncontrolled expansion of “selfish” cellular clones within the host micro-environment^4–7^.

Over the past century, scientists have developed increasingly sophisticated strategies to trace somatic evolution^8–10^ – collectively known as lineage tracing (LT) methods. While foundational, early LT approaches offered limited resolution and phenotypic read-outs, and were largely restricted to experimental models^9,11^. The advent of single-cell multi-omics has since revolutionized the field, enabling the development of single-cell lineage tracing (scLT) technologies^10,12,13^. By jointly profiling cell lineage - via naturally occurring (epi)mutations^14–17^ or exogenous genetic labels^18–26^ - and cell state - including gene expression and chromatin accessibility - at single-cell resolution, scLT methods allow high-resolution mapping of cell fate dynamics across time and space. When applied to both engineered systems and native biological contexts, these approaches have uncovered fundamental principles of tissue development^27–29^ and cancer evolution^30–34^.

Despite their transformative potential, current scLT systems face fundamental trade-offs. While the optimal LT marker should exhibit a fast mutation rate, be cost-effective to sequence, seamlessly integrate with multi-omic workflows, and originate endogenously (e.g., naturally occurring (epi-)genetic mutations), available LT markers struggle to meet all of these requirements. In this scenario, somatic mitochondrial DNA (mtDNA) mutations (MT-SNVs) emerged as a unique alternative due to mtDNA fast mutation rate (10-100x higher than nuclear genome), compact size (∼16 kB), and high sequencing efficiency^14,35,36^. Indeed, several single-cell platforms have been recently developed for simultaneous profiling of MT-SNVs alongside other molecular features^14,37–42^, and analysis of resulting mitochondrial scLT (MT-scLT) datasets revealed key molecular determinants of human hematopoiesis^42^ and Acute Myeloid Leukemia^38,43–45^ clonal evolution.

Despite promising applications, recent contrasting evidence sparked debate over best practices for MT-scLT data analysis^46–49^. Critical concerns revolve around the resolution and robustness of MT-SNV-based lineage inference. Specifically, which types of somatic events – such as early vs late clonal divergence - can be reliably marked by MT-SNVs, and whether MT-SNVs can support robust single-cell phylogeny inference. As a result, the practical scope and reliability of MT-scLT remain uncertain compared to more established scLT systems^46,47^. Addressing this ambiguity is essential for unlocking the full potential of MT-scLT and establishing it as a reliable tool for dissecting somatic evolution in native biological contexts. To address these challenges, here we present `MiTo`, the first end-to-end framework for robust MT-scLT. `MiTo` integrates a flexible `Nextflow` pipeline for data pre-processing and MT-SNVs-based lineage inference with an interactive `python` package for custom, lineage-informed downstream analyses. This toolkit introduces novel algorithms for MT-SNVs and cell filtering, MT-genotyping and lineage inference. When benchmarked against existing baselines and state-of-the-art methods - using lentiviral barcoding datasets with unprecedented size and clonal complexity - `MiTo` consistently outperformed all the alternatives.

To demonstrate its utility in a biologically and clinically relevant context, we leveraged `MiTo` to investigate the poorly understood clonal evolution and phenotypic plasticity of breast cancer (BC). Applied to a longitudinal BC dataset *in vivo*, `MiTo`: i) accurately inferred ground-truth clonal structures and cell state transitions, ii) identified candidate transcriptional determinants of clonal fitness, and iii) quantified heritability of gene expression programs, providing mechanistic insights into tumor progression. To place these results within the broader landscape of single-cell lineage tracing, we used MiTo to compare single-cell phylogenies reconstructed from four evolving scLT markers: MT-SNVs profiled with two distinct experimental protocols (MAESTER^40^ and RedeeM^42^), and nuclear scLT markers, including nuclear SNVs^50^ and Cas9-based evolving barcodes^31^.

This work delivers the first end-to-end solution for MT-scLT, as well the first experimental demonstration that MT-scLT can be used to longitudinally track the phenotypic evolution of ground-truth cell lineages *in vivo*. Beyond introducing a robust analytical framework, our systematic benchmarks establish best-practices for MT-scLT data analysis, expose key strengths and limitations of current scLT systems, and define a roadmap for future methodological innovation in the field.

## Results

To enable robust and scalable mitochondrial scLT (MT-scLT), we developed MiTo - a user-friendly toolkit comprising the `nf-MiTo` pipeline and the `MiTo` package (Fig. 1). We will use the term “phylogeny” to indicate a binary genealogical tree representing the approximate history of cell divisions within a population - with profiled cells as leaves and putative ancestors as internal nodes - and the term “clone” to indicate a genetically defined cell population sharing a recent common ancestor.

**Fig.1.**
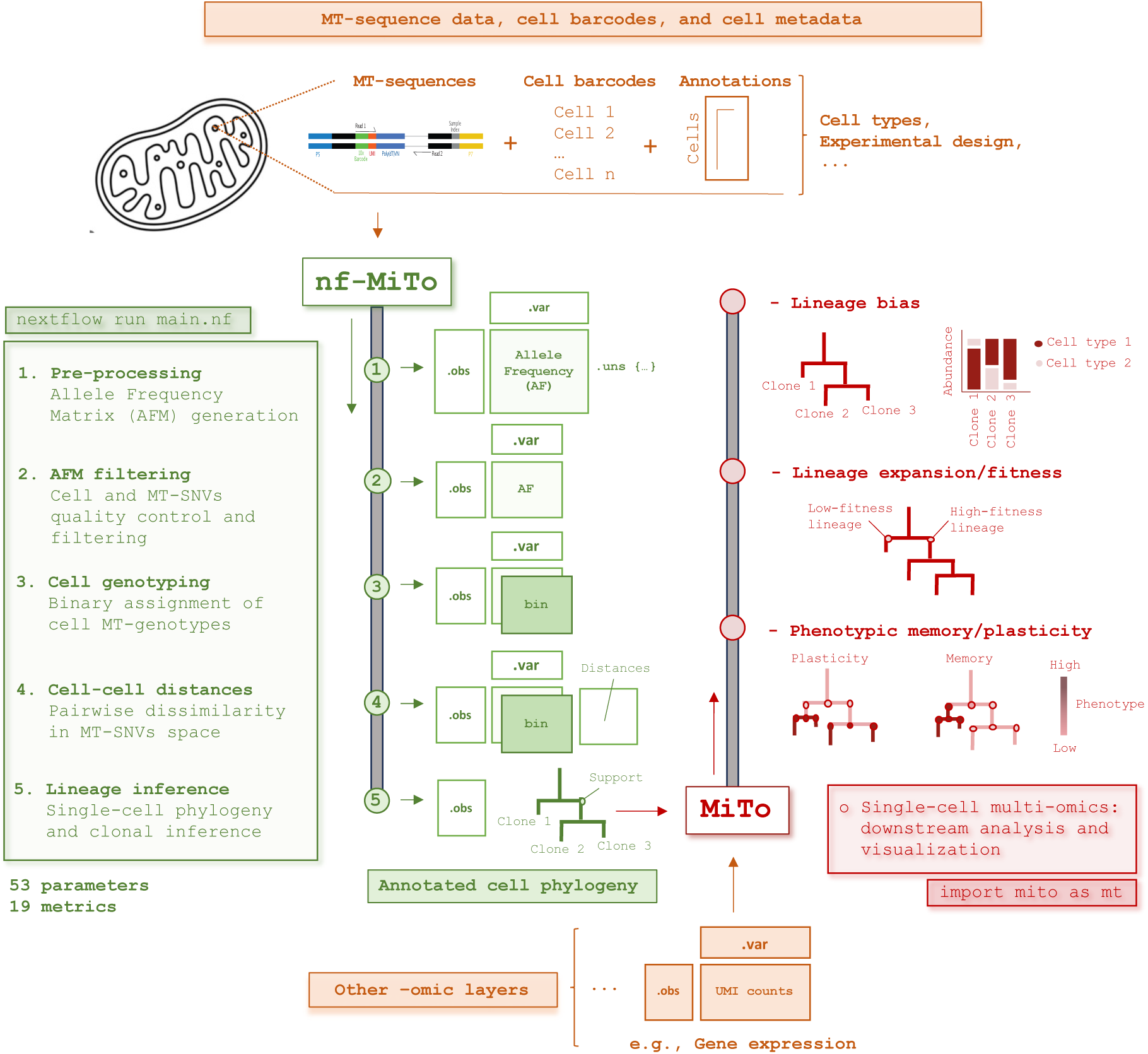
Overview of the MiTo toolkit. Schematic representation of input data (top and bottom, orange) the `nf-MiTo` pipeline (green, left) and the companion `MiTo` package (red, right). The data processing workflow is presented vertically. The top section (orange) illustrates `nf-MiTo` input data. From left to right, these data include raw MT-reads (MAESTER protocol), a list of cell barcodes, and the associated cell metadata (cell type annotations and/or experimental design covariates). The left section (green) shows the main data processing steps of the `nf-MiTo` pipeline – considering default entry-point and hyper-parameters. Step 1: Raw MT reads are processed with `maegatk` to generate an Allele Frequency Matrix (AFM) in `AnnData` format (.X, .var and .obs slots are shown). Cell-level metadata can be added at this stage. Step 2: `MiTo` AFM filters are applied to discard low-quality cells and MT-SNVs. Step 3: MT-genotyping is performed with the `MiTo` probabilistic model, yielding a binarized cell-by-variant matrix (i.e., the bin layer). Step 4: MT-genotypes are used to calculate pairwise cell-cell distances in MT-SNV space (with a custom, weighted Jaccard distance). Step 5: cell-cell distances are used to infer a single-cell phylogeny with the UPGMA algorithm. The resulting tree is subsequently bootstrapped and annotated with intenal node supports and clone-level information (e.g., Clone 1, 2, 3). Users may bypass raw-reads pre-processing (Step 1) by providing directly a pre-processed AFM. The **r**ight section (red) represents the 3 key downstream applications enabled by `MiTo`, starting from an annotated cell phylogeny (`nf-MiTo` output) and matched multi-omic data (bottom section, orange): i) quantification of lineage bias (top, e.g., differential abundance of cell types across MT-clones). ii) integrated analysis of clonal fitness (middle); and iii) quantification of phenotypic hieritability, to dissect memory/plasticity cell phenotypes (bottom). For default parameters and method details, see *Computational Methods* in *Materials and Methods*; for implementation details and alternative `MiTo` entry-points, see *Supplementary Information*. Color code: orange, input data; green, `nf-MiTo`; red, `MiTo`.

`nf-MiTo` leverages `Nextflow` to streamline the five core steps of MT-scLT, including: i) pre-processing of raw MT-reads into an Allele Frequency Matrix (AFM, a 𝑐𝑒𝑙𝑙 𝑥 𝑣𝑎𝑟𝑖𝑎𝑛𝑡 matrix storing the Allelic Frequency of 𝑣𝑎𝑟𝑖𝑎𝑛𝑡_*j*_ in 𝑐𝑒𝑙𝑙_*i*_); ii) filtering high-quality cells and MT-SNVs; iii) assigning binary cell-genotypes; iv) computing pairwise cell-cell distances in the MT-SNV space; and v) inferring lineages, in terms of mitochondrial phylogenies and clones (Fig. 1, left section; Computational Methods in Materials and Methods). Of note, `nf-MiTo` supports input at multiple pre-processing stages - from raw sequencing reads to fully annotated cell phylogenies – allowing users to explore MT-data in automated and scalable fashion (Fig. 1, top section; `MiTo` entrypoints in Supplementary Information).

As the underlying engine of its companion pipeline, `MiTo` not only implements the core functionalities of `nf-MiTo,` but also serves as a standalone `python` package for interactive, lineage-informed, single-cell multi-omic analyses (Fig. 1, right section). Developed according to `scverse` standards^51^ (e.g., the `AnnData` class usage), `MiTo` seamlessly integrates MT-data analysis into widely adopted single-cell workflows (Computational Methods in Materials and Methods). Provided with externally processed single-cell multi-omic data (e.g., gene expression, chromatin accessibility) and metadata (e.g., cell type, donor, tissue of origin), `MiTo` enables interactive analyses of annotated MT-phylogenies from `nf-MiTo`. These downstream analyses include: i) identification of relevant biases in clonal behavior – e.g., uneven cell type composition across MT-clones; ii) prioritization of molecular features significantly associated with clonal fitness; iii) quantitative measurement of cell phenotype heritability - including gene expression, chromatin states, and other experimental covariates – to distinguish phenotypic plasticity from memory (Fig. 1, right section).

To date, `nf-MiTo` and `MiTo` together represent the first end-to-end solution for MT-scLT. This toolkit advances MT-scLT state-of-the-art by integrating newly developed methods – including optimized algorithms for MT-reads pre-processing, AFM filtering, MT-genotyping, and clonal inference – with established tools into a unified, modular, and extensible framework. For a general overview, see Computational Methods in Materials and Methods. For a detailed breakdown of `nf-MiTo` and `MiTo` capabilities, see `MiTo` toolkit in Supplementary Information.

### New benchmarking datasets with ground-truth lineages

To systematically benchmark our toolkit, we required datasets with known (ground-truth) lineage relationships. Existing strategies in the field include both computational simulations ^49^ and experimental cell labelling^14,42^. Although simulations allow precise control over data generation, unrealistic assumptions and/or parameter values might produce overly simplified benchmarking datasets. However, existing real-world benchmarking datasets fail to capture the cellular and clonal complexity of current MT-scLT experiments. Among the first, Ludwig et al.^14^ generated two benchmarking datasets combining measurement of expressed MT-SNVs and lentiviral barcodes. However, these datasets included only 3-11 ground-truth lentiviral clones and 70-158 cells, respectively. More recently, Weng et al.^42^, introduced ten new benchmarking datasets combining measurement of MT-SNVs and Cas9-based evolving barcodes. However, even in this case, the number of ground-truth clones and cells per dataset remain limited (<10 clones and 200-600 cells, respectively). To generate benchmarking dataset that more faithfully reflect the scale and complexity of current MT-scLT experiments, here we performed two dual-lineage tracing experiments combining measurement of expressed MT-SNVs and lentiviral barcodes (Supp. Fig. 1; Data Generation in Materials and Methods).

In the first experiment, individual cells from the breast cancer MDA-MB-231 cell line were isolated by FACS and expanded *in vitro* to generate single-cell derived colonies. After 30 days, each colony was transduced with a unique barcode-expressing lentivirus. Then, following a short selection period (∼7 days), eight barcoded colonies were mixed in defined proportions to create a controlled clonal mixture (MDA_clones).

In the second experiment, MDA-MB-231 cells transduced with the high-complexity Perturb-seq lentiviral barcode library^52^ were orthotopically transplanted into immunocompromised mice. ∼30 days post-injection, primary tumors (PTs) were surgically resected, and, after additional ∼30 days, mice were sacrificed and lung metastases collected. One matched PT-lung pair was selected for further data generation (MDA_PT and MDA_lung, respectively).

Cell suspensions from all three specimens underwent single-cell RNA sequencing (scRNAseq). To enable simultaneous recovery of gene expression (GEX), mitochondrial variants (MT), and expressed lentiviral barcodes (GBC), we modified the original MAESTER protocol^40^ to include the additional GBC read-out (Supp. Fig. 1; Data Generation in Materials and Methods and Supplementary Information). After raw data pre-processing and stringent Quality Control (Materials and Methods – Data Analysis) we obtained three high-quality multi-modal datasets (i.e., MDA_clones, MDA_PT, and MDA_lung datasets) with variable numbers of GBC clones (n=8-216; Fig. 2a, left panel), cells (n=375-2757; Fig. 2a, center panel), GBC clonal complexity (Shannon Entropy = 0.77-1.69; Fig. 2a, right panel, and Fig. 2b) and MT-genome coverage (n consensus UMIs = 44-83; Fig.2c).

**Fig.2.**
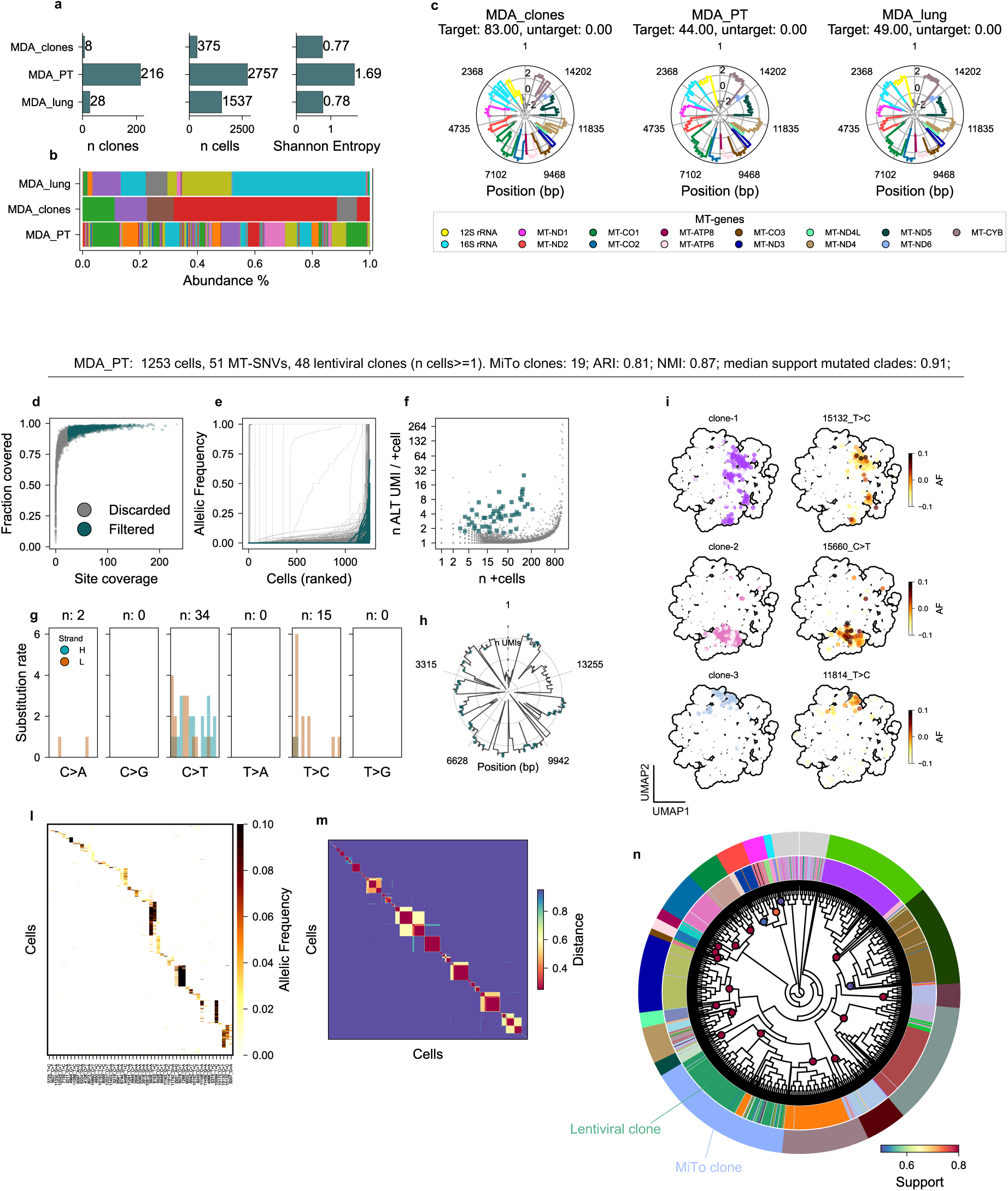
Benchmarking datasets and representative output from nf-MiTo pipeline. **a.** Bar plots showing summary statistics for each benchmarking dataset: number of ground-truth clones, number of cells, and clonal complexity (measured by Shannon Entropy). All statistics are computed after Cell QC, and *before* additional AFM filtering. See Cell QC in Materials and Methods **b.** Stacked bar plots representing clonal prevalences across the three benchmarking datasets. Color-coding refers to lentiviral clone identity. **c,** Mitochondrial coverage plots, for each dataset. Each plot shows the mean coverage (radial axis) across the mitochondrial genome (1–16,569 bp, anti-clockwise). Consensus UMIs (from `maegatk`, with minimum 3 supporting reads) are counted. Coverage annotations refers to targeted and untargeted regions from the MAESTER protocol. Color-coding indicate MAESTER-targeted mitochondrial genes. **d.** `MiTo` cell filtering, based on mitochondrial genome coverage statistics. Each dot represents a cell. x-axis: mean site coverage across the MT-genome. y-axis: fraction of MAESTER-targeted sites covered (n consensus UMIs>0). See Computational Methods in Materials and Methods. Color-coding indicated filtered and discarded cells (green and grey, respectively). **e.** MT-SNV filtering. Each line shows the ordered vector of Allelic Frequencies (AF) of a single MT-SNV across all cells. Color-code is similar to d, but for MT-SNVs instead of cells. **f.** MT-SNV filtering. Each point represents a MT-SNV. x-axis: number of positive (AF>0) cells. y-axis: mean number of alternative (ALT) consensus UMIs in positive (AF>0) cells. Color-code as in e. **g.** Mutational spectrum of filtered MT-SNVs, enriched for T>C and C>T substitutions (H: heavy strand; L: light strand on the MT-genome). **h.** Mitochondrial genome coverage plots (as in c) with filtered MT-SNVs (green crosses), showing their genomic distribution. **i.** UMAP plots highlighting selected GBC clones (left column) and their corresponding enriched MT-SNVs (right column). UMAP coordinates embed embed cells into 2D space, with distances reflecting dissimilarity in the MT-SNV space. **l.** Filtered Allele Frequency Matrix (AFM) shown as a clustered heatmap (n cells = 1,253, n MT-SNVs = 51). **m.** Clustered weighted Jaccard distance matrix (order as in **n**). See Computational Methods in Materials and Methods. **n.** MT-phylogeny – UPMGA algorithm - depicted as an annotated dendrogram. Internal and external color strips indicate GBC and MT clones, respectivel. Light grey indicates unassigned cells (<10% total cells). Colored nodes represent clonal nodes, annotated with their Transfer Bootstrap Support. See Computational Methods in Materials and Methods for details.

Collectively, these 3 high-quality datasets substantially exceeded previous ones in terms of both number of cells and ground-truth clones, representing three real-world scenarios of increasing complexity: i) low cells and clone counts, with balanced clonal frequencies (MDA_clones); ii) high cell but low clone counts, with unbalanced clonal frequencies (MDA_lung); and iii) high cell and clone counts, with relatively-balanced clonal frequencies (MDA_PT).

### MiTo benchmark

Next, we leveraged our three benchmarking datasets to systematically tune `nf-MiTo` hyper-parameters - see `nf-MiTo` hyper-parameters in Supplementary Information.

Although lineage inference occurs only at the final step of the `nf-MiTo` pipeline (Fig.1, step v), its output depends on all upstream processes (Fig.1, step i-iv) and their associated hyper-parameters. Thus, we adopted a local optimization strategy. First, we set a baseline configuration for `nf-MiTo` for hyper-parameters (Supp. Fig. 2; see also Preliminary assessment in Supplementary Information), and then we performed a step-wise ablation analysis. Specifically, we perturbed one or more hyper-parameters at the time, assessed their impact on lineage inference performance - the concordance between inferred and ground-truth clones balanced by the number of cells and ground-truth clones retained after MT-data pre-processing (Fig.1, step i-iii, see `MiTo` benchmark in Supplementary Information) –, and changed baseline values accordingly. Multiple iterations of this process refined `nf-MiTo` baseline configuration, quantified the performance gains introduced by our framework (see Computational Methods in Materials and Methods) and revealed previously underappreciated steps of the MT-scLT workflow (see `MiTo` benchmark in Supplementary Information).

In particular, robust MT-data pre-processing proved essential for accurate lineage inference. Among raw read processing strategies, UMI-based consensus error correction – specifically `maegatk` and `MiTo` – was the only approach that enabled accurate clonal inference across all datasets (Adjusted Rand Index (ARI) >0.8, see Supp. Fig. 4-7). However, this accuracy was strictly dependent on `MiTo` cell and MT-SNVs filtering. To filter only cells with high-quality MT-libraries, `MiTo` leverages MT-genome coverage metrics (Fig. 2d and Supp. Fig. 3, see Computational Methods in Materials and Methods). Despite substantial loss of cells (20-60% of cells passing gene-expression quality control, see Cell QC in Materials and Methods), our data showed that more relaxed cell filtering dramatically impairs clonal inference accuracy (Supp. Fig. 2-3), underscoring the importance of this step. To filter high-confidence MT-SNVs, instead, `MiTo` integrates variant-level summary statistics with external databases and custom statistical tests to remove noisy or un-informative variants (Fig. 2e-l and Supp. Fig. 7-8, see also Computational Methods in Materials and Methods). Importantly, each filtering criterion effectively improved high-confidence MT-SNVs detection and lineage inference performance (Fig. 2i-l). Provided with high-quality cells and high-confidence MT-SNVs, `MiTo` probabilistic framework for MT-genotyping (Fig. 1, step iii) outperformed simpler baselines (Supp. Fig. 11-12; see MT-genotyping in Materials and Methods and Supplementary Information), especially under conditions of artificially introduced noise (Supp. Fig. 13). Taken together, our data showed that `maegatk` raw reads pre-processing combined with `MiTo` AFM filtering and genotyping constitutes the best pre-processing workflow for expressed MT-scLT – i.e., the workflow achieving the best trade-off between clonal inference accuracy and cellular yield (Supp Fig. 2 and 9).

Provided with robustly pre-processed MT-data and cell-cell distances (Fig.1, step i-iv), distance-based tree-building algorithms – such as Neighbor Joining and UPMGA - inferred MT-phylogenies (Fig.1, step v) that hierarchically clustered ground-truth clones, as shown for the representative MDA_PT dataset in Fig. 2m-n. To test whether cutting these MT-phylogenies into MT-SNVs-supported clades (Fig. 2n) yields more accurate and finely resolved clones than alternative heuristics, we benchmarked `MiTo` clonal inference against 3 state-of-the-art methods: i) `leiden`^53^, a graph-based community detection algorithm; ii) `vireoSNP`^54^, a Bayesian clustering method; and iii) `CClone`^55^, a Non-Negative Matrix Factorization (NMF) -based algorithm (see Clonal inference benchmark in Materials and Methods). In particular, we selected 15 AFMs with strong phylogenetic signal – five alternative selections of cells and MT-SNVs for dataset (see associated statistics in Fig. 3a) and performed lineage inference optimizing each method hyper-parameters (as detailed in Clonal inference benchmark, Supplementary Information). As shown in Fig. 3b-c, `MiTo` consistently outperformed other methods across all datasets, considering both the number of inferred clones – compared to the ground truth, Fig. 3b - and the concordance between inferred and ground-truth clonal labels (Fig. 3c). Compared to `MiTo`, `vireoSNP` – the 2^nd^ ranked method - achieved comparable accuracy and resolution in low-complexity datasets (i.e., MDA_clones, MDA_lung), but significantly lower resolution in the high-complexity dataset (0.5x number of ground-truth clones detected in MDA_PT). `MiTo` run time also scaled better with dataset size, while memory requirements were substantially low for both methods - i.e., both can be safely run on a modern laptop with 8GB and 16 cores (Supp. Fig. 15).

**Fig.3.**
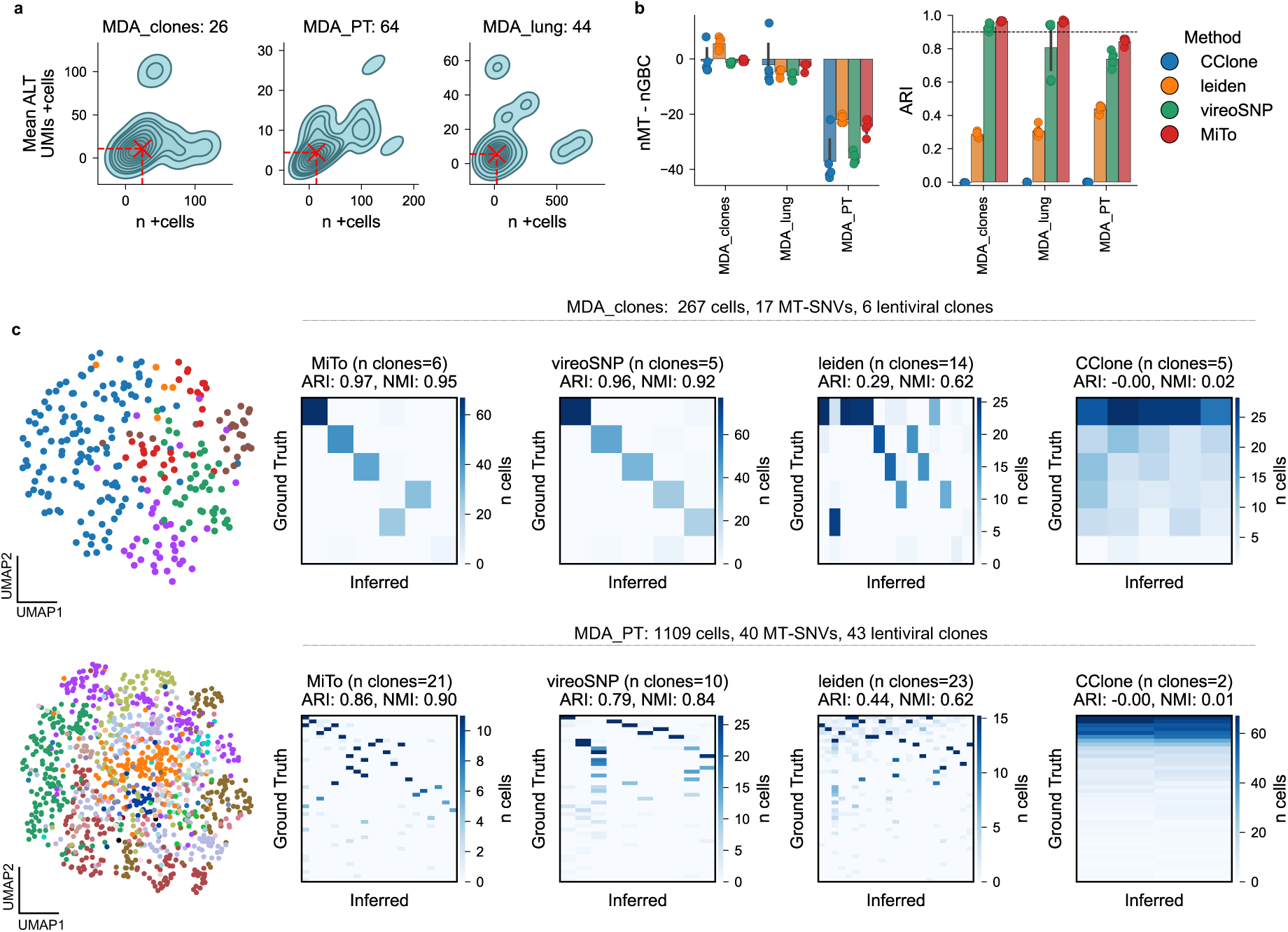
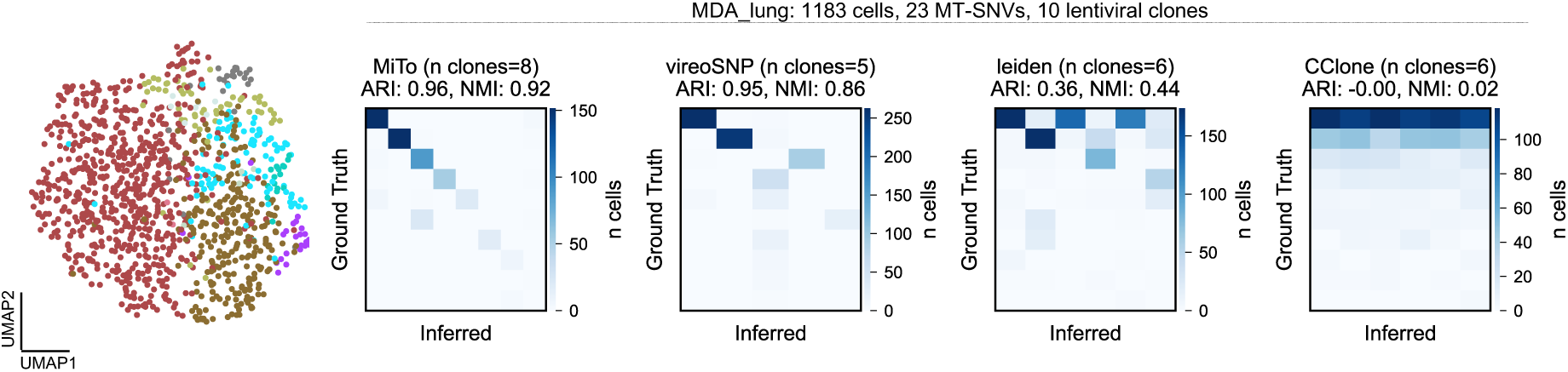
MiTo clonal inference benchmarked against 3 state-of-the-art methods. **a.** 2-dimensional density plots showing the distribution of 2 key summary statistics across filtered MT-SNVs: the number of positive cells (x-axis); and the mean number of alternative (ALT) UMIs in positive cells (y-axis, see also Fig. 2f). Each point is a filtered MT-SNVs. Each panel is annotated with the total number of filtered MT-SNVs for the respective dataset – i.e., the union of filtered MT-SNV spaces (n=5 per dataset) used for the clonal inference benchmark. Dashed red lines indicate x- and y-axis medians. **b.** Clonal inference performance summary. Left: difference between the number of inferred clones (n_MT) and ground-truth clones (n_GBC). Right: Adjusted Rand index (ARI) between inferred and ground-truth clonal labels. Each dot represents a distinct inference task (n=5 per dataset). **c.** Detailed visualization of representative clonal inference results. One representative MT-SNV space per dataset (row) is shown (i.e., the MT-SNV space with the highest mean ARI across clonal inference methods). Leftmost panels: UMAP - representing cell-cell dissimilarity in MT-SNV space - color-coded by ground-truth GBC clone. Remaining columns: confusion matrices comparing ground-truth (GBC clones, heatmap rows) and inferred (MT-clones, heatmap columns) clones, for each method. Clonal inference metrics (ARI, NMI) and number of inferred clones are annotated above each matrix.

These data demonstrate that `MiTo` yields more accurate and resolved clonal inferences than alternative methods, providing also informative ancestor-descendant relationships between inferred mitochondrial (MT-) clones.

Together, our systematic benchmark validates `MiTo` and `nf-MiTo` as robust and flexible solution for MT-scLT, demonstrating accurate inference of ground-truth cell lineages from benchmarking datasets of unprecedented size and clonal complexity. These results lay the foundation for broad applications across diverse biological and clinical contexts.

### MiTo traces the phenotypic evolution of breast cancer clones *in vivo*

To assess the performance of `MiTo` in a biologically and clinically relevant context, we applied it to our longitudinal PT-lung dataset (Fig. 4a). We first analyzed gene expression and lineage – GBC and MT – independently, performing cell state annotation and lineage inference on the combined MDA_PT and MDA_lung dataset (see Multi-omic analysis of longitudinal Breast Cancer clones in Materials and Methods). The final multi-modal dataset included 2 time-points, 2549 cells, 77 MT-SNVs, 13057 expressed genes, 33 MT- and 60 GBC-clones. Notably, MT and GBC clones showed very high concordance even across time-points (ARI=0.94, Fig, 4c and Supp. Fig. 16-17), demonstrating robustness of `MiTo` clones. Integration of lineage and gene expression modalities revealed a continuous trajectory from PT to lung metastasis, marked by a gradual shift in dominant cell states (Fig. 4b–c). PT cells were enriched for TGFβ– EMT, IFNα/γ, Hypoxia, and Proliferation, whereas lung metastasis predominantly exhibited an OXPHOS phenotype, consistent with metabolic adaptation during metastatic progression.

**Fig.4.**
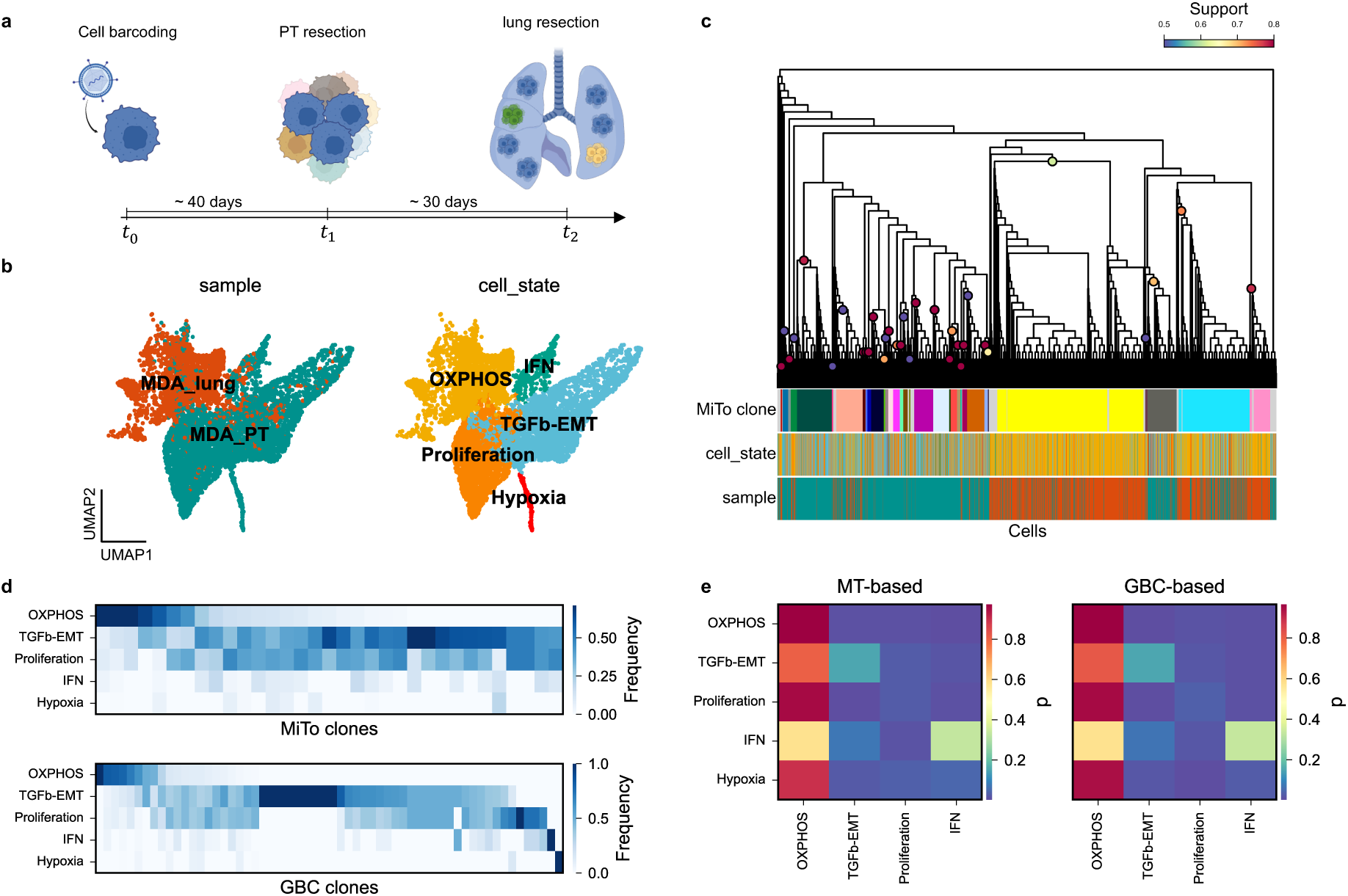
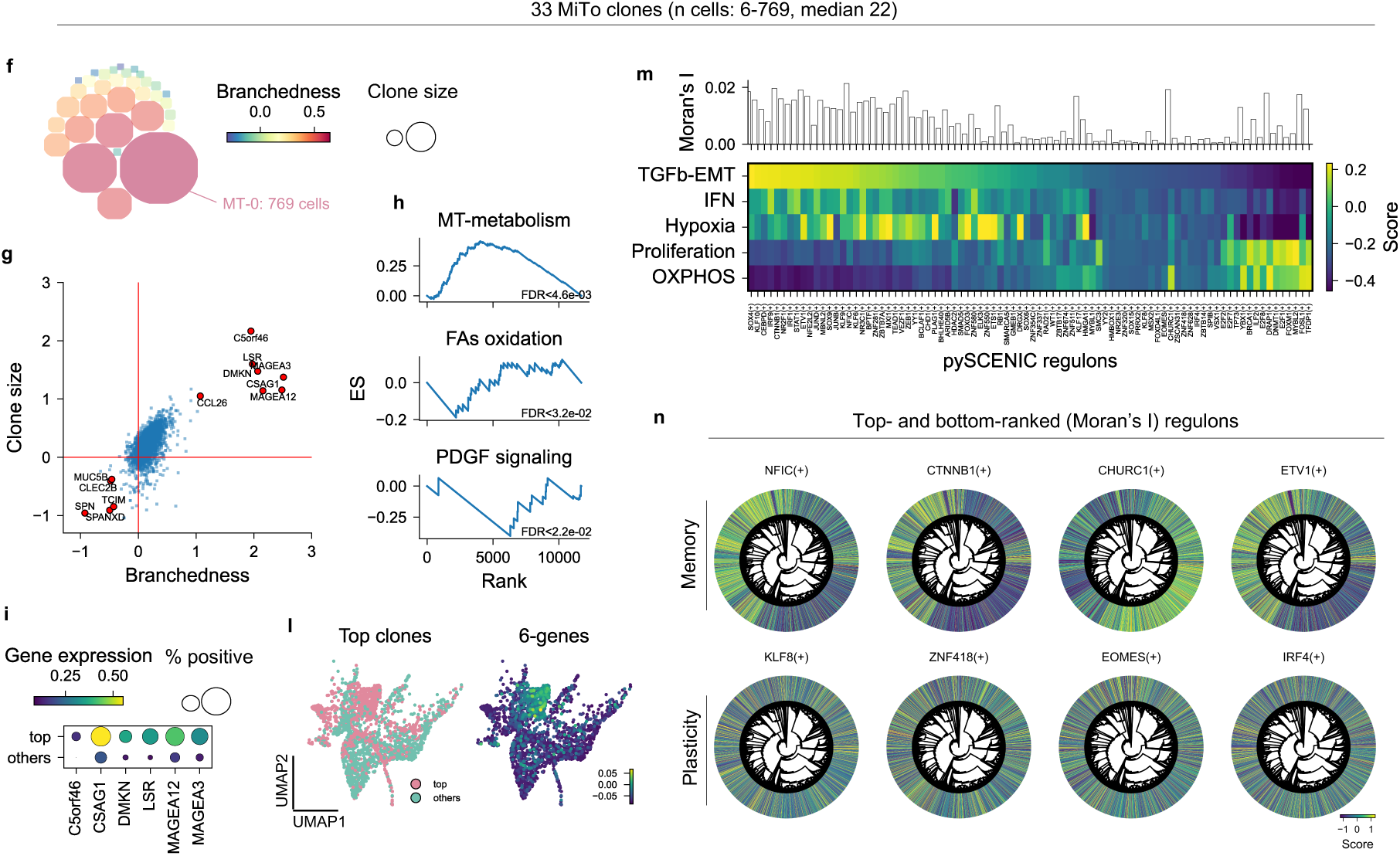
MiTo trace the phenotypic evolution of breast cancer clones *in vivo.* **a.** Schematic representation of the experimental design used to generate the longitudinal PT-lung dataset (see Data generation in Materials and Methods). **b.** UMAP plots of the combined PT-lung dataset. (n = 2,549 cells). Left: cell colored by tissue of origin (PT or lung). Right: cells colored by cell state (legends on plots). See Multi-omic analysis of longitudinal Breast Cancer clones in Materials and Methods. **c.** Dendrogram showing the MT-phylogeny of the PT-lung dataset (UPMGA algorithm; n=77 MT-SNVs, n=2549 cells). Colorstrips annotate each cell tissue of origin and cell state (color code as in panel c). **d.** Heatmaps showing cell state distributions of MT-clones (top) and GBC-clones (bottom). Each row represents a MT- or GBC-clone. Values are row-normalized (i.e., each row sums to 1). **e.** Cell state transition probabilities computed with Moslin using either MT-based (left) or GBC-based (right) cell–cell distances (See Multi-omic analysis of longitudinal Breast Cancer clones in Materials and Methods). Each 𝑖𝑗 entry represent the (average) probability of a cell in state 𝑖 in the PT to reach cell state 𝑗 in the lung. **f.** Circle-packed plot representing the size (area) and branchedness (color) of each MT-clone. The brancheness metric measures the topological complexity of each clone’s sub-tree on the MT-phylogeny. **g.** Volcano plot of gene-wise negative binomial regression coefficients for clone branchedness (x-axis) and size (y-axis). Each dot represents a gene. Selected genes are annotated (see Multi-omic analysis of longitudinal Breast Cancer clones in Materials and Methods). **h.** Gene Set Enrichment Analysis (GSEA; Methods) results for 3 GO terms and MsigDB pathways. For each panel, the x-axis represents the input gene list rank, and the y-axis represents the GSEA Enrichment Score (ES, see Methods). **i.** Dot plot showing mean gene expression (color) and fraction of expressing (n UMIs>0) cells (size) for the six fitness-associated genes shown also in panel g. Single-cell gene expression values are aggregated (mean) for either the top3 size-ranked clones, and all the others cells. **l.** UMAP plots showing top3 clones’ cells and the 6 genes signature score (see Methods). **m.** pySCENIC regulons activation in cell states. Each heatmap value represent the mean activation score (Methods) for regulon 𝑗, (column) in cell state 𝑖 (row). The barplot on the upper panel show Moran’s I statistic (Methods) computed for each regulon using distances in MT-SNVs and regulon activation scores as inputs. **n.** Dendrograms representing the same MT-phylogeny in **b,** in polar coordinates. Colorstrips show regulon activation scores for each cell (tree leaves).

Comparison of GBC and MT-clones revealed similar cell-state biases: high-prevalence clones from both modalities enriched for the OXPHOS state, while intermediate-to-low prevalence clones enriched for the TGFβ-EMT and proliferative states. One notable exception was represented by two low-prevalence GBC clones enriched for the hypoxia and IFN cell states that were not detected as distinct MT-clones (Fig 4d). To quantitatively validate these trends, we computed cell state transition probabilities using `Moslin`^56^ providing either GBC-or MT-pairwise cell-cell distances (see Clonal bias and cell state transition dynamics in Materials and Methods). Strikingly, GBC- and MT-estimated cell state transitions exhibited strong qualitative and quantitative agreement, demonstrating how GBC and MT lineage modalities can be equivalently used to measure dynamic cellular processes (Fig 4e).

Next, we associated quantitative descriptors of clonal fitness with gene expression data (see -Fig. 4 f-l and Transcriptional determinants of clonal fitness in Materials and Methods). Here, we found that increased clonal fitness correlates increased mitochondrial metabolism, reduced Fatty-Acid Oxidation, and reduced Platelet-Derived Growth Factor (PDGF) signaling (Fig. 4h). We also identified six genes significantly up-regulated in high-fitness clones (see Fig. 4 g, i, l, and Transcriptional determinants of clonal fitness in Materials and Methods): C5orf46, CSAG1, DMKN, LSR, MAGEA12, and MAGEA3. Importantly, several of these genes (e.g., MAGEA3, LSR) have established roles in cancer progression^57–59^, while others (e.g., C5orf46, DMKN) have been implicated as markers of poor clinical prognosis^60–62^. These findings highlight how incorporating lineage-derived fitness descriptors enables biologically meaningful marker gene discovery that may be overlooked by alternative approaches.

Finally, to investigate heritability of gene regulatory programs, we inferred gene regulatory networks with `pySCENIC`^63^ and quantified their phylogenetic autocorrelation across the MT-phylogeny using Moran’s I statistic (Fig. 4m-n; see Memory and plasticity of gene expression programs in Materials and Methods). Among 81 regulons differentially activated across cell states (Fig. 4m), most showed moderate yet statistically significant Moran’s I value. These data suggest that a substantial subset of transcriptional programs retains lineage “memory”, reflecting heritable regulatory constraints during tumor evolution.

Mapping regulon activity onto the MT-phylogeny (Fig. 4n) allowed us to visualize “memory” and “plasticity” regulons, defined by either clustered or widespread activation patterns across the MT-phylogeny. Of note, memory regulons were regulated by transcription factors (TFs) such as NFIC, CTNNB, CHURCH1 and ETV1, previously associated with the maintenance of epithelial cell identity. Conversely, plasticity regulons were driven by KLF8, ZNF418, EOMES, and IRF4 TFs, linked with epithelial-to-mesenchymal (EMT) transition, stemness and stress response. These patterns suggest that cancer cell identity and behavior is shaped by distinct layers of transcriptional regulation, where both lineage-imprinted (e.g., NFIC, CTNNB, CHURCH1) and environment-responsive programs (e.g., KLF8, EOMES, IRF4) co-exist to drive tumor progression.

Together, these findings highlight `MiTo` unique ability to extract biological insights from MT-scLT data. By accurately reconstructing lineage relationships, quantifying clonal dynamics, and linking them to transcriptional phenotypes, `MiTo` provides a powerful framework to dissect the molecular determinants of somatic evolution *in vivo*.

### MT-scLT in the broader scLT landscape: limitations and opportunities

Several studies have demonstrated the practical utility of MT-scLT in diverse biological and clinical contexts^41,42,44,45^. However, recent conflicting reports have raised concerns about the resolution and reliability of mitochondrial phylogenies (MT-phylogenies)^46,47,49^. In addition, a systematic comparison between single-cell phylogenies inferred from MT-SNVs – called from either MT-DNA or RNA – and other nuclear LT markers is still lacking.

To address this gap, we leveraged `nf-MiTo` to compare single-cell cell phylogenies across four evolving scLT systems: i) single-colony whole-genome sequencing (scWGS)^50^; ii) Cas9-based molecular recording^31^; iii) the RedeeM protocol^42^, enriching for MT-DNA molecules in 10x Multiome libraries; and iv) the MAESTER protocol^40^, as implemented in this study. We analyzed n=16 datasets (13 public and 3 from this work, see Fig. 5a), including 3 scWGS datasets, 5 Cas9 datasets, and 5 RedeeM datasets (Fig. 5a). For each of these datasets we collected pre-processed character matrices and metadata, and then used `nf-MiTo` to infer single-cell phylogenies and evaluate their properties metrics (Fig. 5b-c, see `nf-MiTo` metrics in Supplementary Information). This comparative analysis provided a quantitative basis to evaluate the robustness and resolution of MT-scLT - compared to other scLT systems.

**Fig.5.**
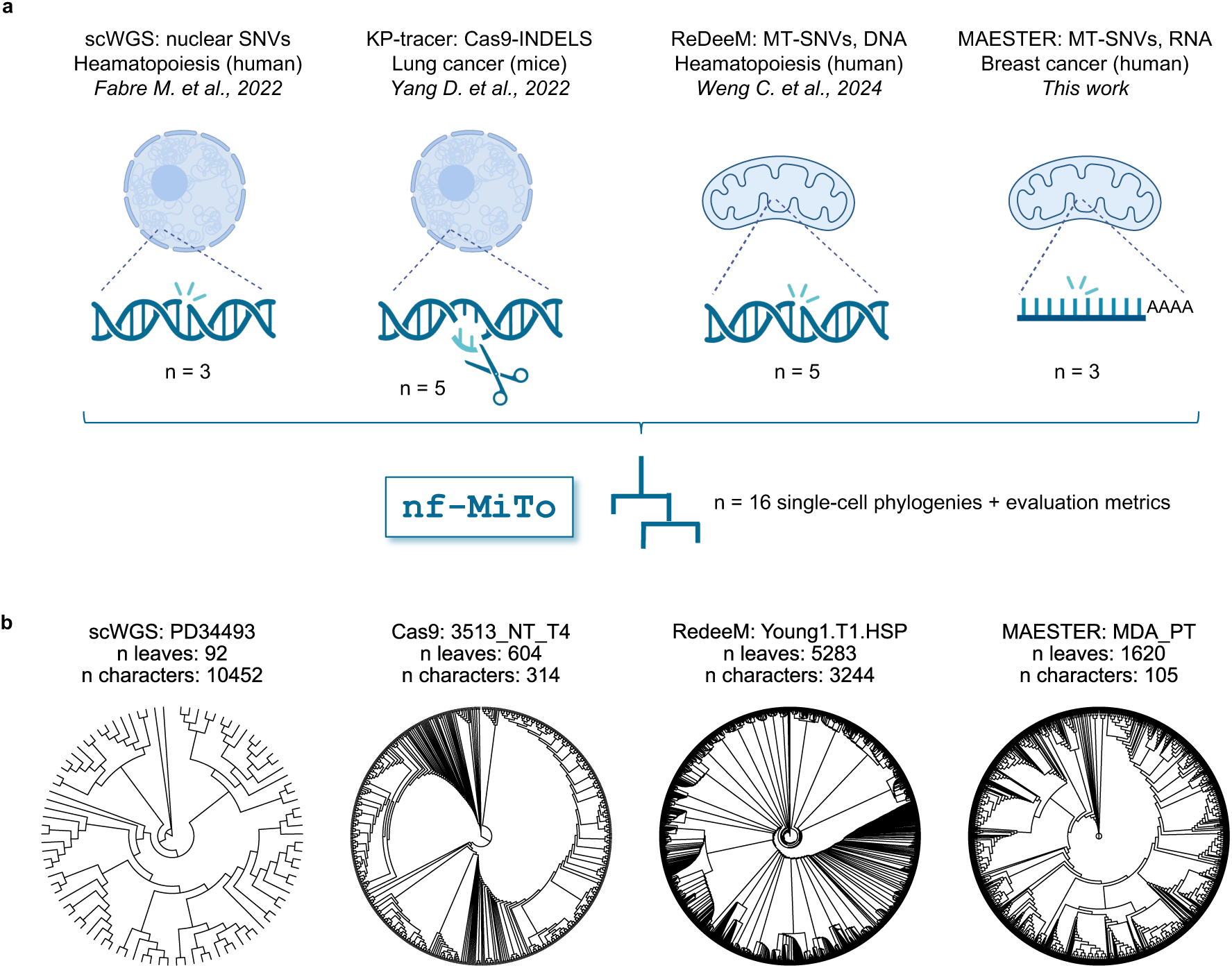
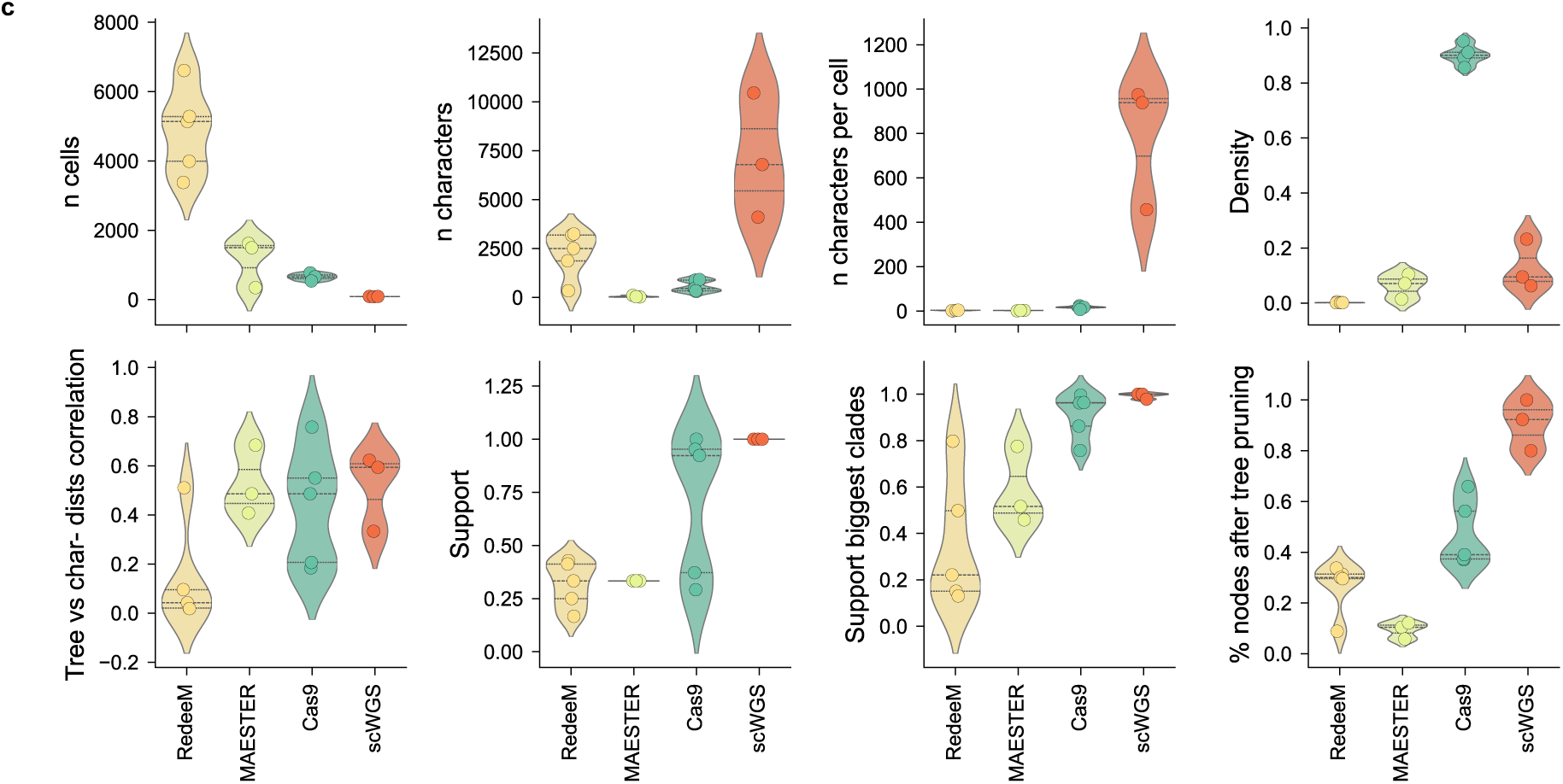
Systematic comparison of single-cell phylogenies across four dynamic scLT systems. **a.** Cartoon illustrating scLT datasets (re-)analyzed. `nf-MiTo` (INFER entrypoint) was run n=16 datasets: 3 scWGS (somatic nuclear SNVs), 5 Cas9 (Cas9 molecular recorders), 5 RedeeM (MT-SNVs from MT-DNA), and 3 MAESTER (MT-SNVs from MT-RNA) datasets. Annotations refers to the original works from which pre-processed data were collected and re-analyzed. **b.** Representative dendrograms for each scLT system phylogenies. For each phylogeny, lineage marker, sample name, number of cells and characters are annotated. **c.** Violin plots with selected `nf-MiTo` metrics grouped by scLT system. Each point represents one dataset. Top row: *n cells* - number of cells (tree leaves) per dataset; *n characters* - total number of characters used for phylogeny inference; *n characters per cell* – number of characters per cell; *Density*: fraction of character matrices non-zero entries. Bottom row: *Tree vs character distance correlation* - correlation between pairwise tree-based and character-based cell-cell distances, reflecting signal-to-noise ratio; *Support* - mean bootstrap support across all internal nodes; *Support biggest clades* - bootstrap support for the largest clades only (top 10^th^ percentile); *% nodes after tree pruning* - fraction of internal nodes retained after mutationless edges have been pruned from the phylogeny.

Our analysis revealed substantial differences among reconstructed phylogenies, as summarized by Fig. 5c. scWGS phylogenies, although limited to small cell numbers (n<100), displayed 10-100x more characters per cell than other methods. This exceptional character diversity translated into high robustness to AFM bootstrapping - across all levels of tree depth - and the highest signal-to-noise ratio, as measured by the correlation between character-based and tree-based pairwise distances (i.e., distance correlation, see Fig. 5c and nf`-MiTo` metrics in Supplementary Information). Cas9 phylogenies, comprising hundreds of cells, ranked second in both robustness and signal-to-noise ratio, followed by MT-phylogenies. Considering all levels of tree depth, these large-scale phylogenies – including 10–100x more cells than Cas9 or scWGS ones - showed low bootstrap support (median<0.5). However, higher support values (>0.7) were typically observed for large (top 10 percentile, Fig. 5c) and mutation-assigned clades (Fig. 2m), suggesting that MT-phylogenies can indeed capture robust lineage relationships among cells, albeit at low resolution. Interestingly, we observed similar supports for 2 out of 5 Cas9 phylogenies, despite the absence of mitochondrial-specific sources of noise. Signal-to-noise ratio also varied across MT-scLT systems, as MAESTER phylogenies showed distance correlations comparable to Cas9 and significantly higher than RedeeM ones (Fig. 5c). These differences highlight marked differences in the phylogenetic signal of MAESTER and RedeeM MT-SNV spaces, which might be reduced by further optimization of RedeeM data pre-processing.

Overall, these data confirm scWGS as the most robust system for high-resolution scLT, despite significant drawbacks - high costs, low throughput and limited multi-omic compatibility. While Cas9-based methods offer a better balance between scLT resolution and scalability, they also require complex genetic engineering, and thus are restricted to model systems. On the other hand, MT-scLT approaches are scalable, label-free, and can provide accurate lineage inferences (Fig.2, Fig.3, Supp. Fig. 17), despite limited resolution. Thus, for studies that prioritize broad scalability and phenotyping over ultra-high lineage inference resolution, MT-scLT offers a practical, cost-effective way to investigate somatic evolution in primary tissues. Nevertheless, caution is warranted when interpreting MT-phylogenies, as many internal nodes – especially at high tree depths - may reflect stochastic noise rather than true clonal divergence events.

## Discussion

Somatic evolution underlies key biological processes in multicellular organisms, including tissue development, regeneration, and cancer progression. Yet, deciphering the molecular mechanisms driving these cellular dynamics remains a major challenge in modern biology. Lineage tracing (LT) methods have long served as fundamental tools for studying cell behavior and fate decisions, and with the recent advancements of single-cell technologies, these approaches have achieved unprecedented resolution and phenotyping depth^10,12^. Among emerging single-cell LT (scLT) strategies, mitochondrial scLT (MT-scLT) has recently emerged as scalable and label-free approach for direct reconstruction of lineage hierarchies in human tissues^37,40,42–45^. However, best practices for analyzing MT-scLT data remain unclear, and the specific contexts in which this technology can provide reliable insights are still being actively debated^46–48^.

To address these challenges, we developed `MiTo`, the first comprehensive, end-to-end solution for MT-scLT data analysis. `MiTo` integrates both established tools and novel algorithms within a modular `Nextflow` pipeline and a user-friendly python package. Unlike existing frameworks^40,42^, `MiTo` supports automated analysis of MT-scLT data, from raw sequencing reads to fully annotated MT-phylogenies. Thanks to its extensive suite of hyper-parameters (n=53) and diagnostic metrics (n=19), `MiTo` offers unparalleled control of its entire data processing workflow, enabling highly customizable and fine-tuned analyses. Importantly, `MiTo` adopts the gold-standard data structure for single-cell omics - the `AnnData`^64^ class - ensuring seamless integration with current and future single-cell multi-omic workflows. Its modular DSL2 syntax also facilitates straightforward integration of new features. These key design choices, along with containerized, platform-agnostic, and HPC-compatible execution, guarantee scalable and reproducible usage across institutions and collaborative projects, representing a significant advancement in the field.

To rigorously evaluate `MiTo`’s performance, we generated three real-world benchmarking datasets with experimentally defined ground-truth lineage relationships – obtained through lentiviral barcoding. Compared to previous efforts, these datasets include substantially higher number of cells and ground-truth clones – up to 2,757 quality-controlled cells and 216 ground-truth clones before Allele Frequency matrices filtering -, more accurately reflecting the complexity and scale of current MT-scLT experiments^41,42,45^. Importantly, these publicly available data lay the foundation for further model development and benchmarking, contributing to improve MT-scLT even beyond this work.

We leveraged these data to benchmark `MiTo` against multiple baselines and state-of-the-art methods. Systematic assessments demonstrated `MiTo`’s superior performance across several key tasks, including MT-SNVs and cell filtering, MT-genotyping, and lineage inference. Here, we demonstrate that pre-processing of MT-scLT data is essential for robust lineage inference, and propose a finely-tuned recipe that effectively balance lineage inference accuracy with cellular yield. Furthermore, we demonstrate that single-cell phylogenies inferred from MT-SNVs (MT-phylogenies) can be used to solve discrete cell clones at higher accuracy and resolution compared to alternative methods – i.e., spectral and Bayesian clustering, Non-Negative Matrix Factorization (NMF)-based approaches. This approach also scales better with dataset size, which is expected to grow in the next years.

As a proof-of-concept, we applied `MiTo` to a time-resolved scLT experiment tracing breast cancer clonal evolution *in vivo*. In this context, `MiTo` solved 30 mitochondrial clones that showed extremely high concordance with ground-truth lentiviral labels (n cells=2549; n ground-truth clones=60; ARI=0.94). While previous studies have reported persistent detection of MT-SNVs in longitudinal experiments - both *in vivo* and in primary tissues^43,45^ – they lacked ground-truth clonal labels, preventing rigorous validation of lineage inference accuracy. To our knowledge, this represents the first experimental demonstration of accurate, time-resolved MT-scLT validated against ground-truth clonal identities. Critically, our multi-omic analyses demonstrate that MT-scLT can provide biological insights that are both qualitatively and quantitatively similar to those derived from lentiviral barcoding — including detection of lineage biases and reconstruction of dynamic cell-state transitions — without the need for experimental cell labeling. When exploring lineage-phenotype associations and cell state heritability, `MiTo` generated a rich set of mechanistic hypotheses regarding the molecular drivers of breast cancer progression *in vivo*. For instance, we found that higher-fitness clones exhibit extensive metabolic rewiring and upregulates a six-genes (C5orf46, CSAG1, DMKN, LSR, MAGEA12, and MAGEA3), previously linked to cancer progression of poor prognosis in breast and/or other cancer types. Moreover, we observed that transcriptional programs governing epithelial identity and stress responses showed distinct modes of inheritance across tumor evolution, highlighting the complex interplay between lineage memory and plasticity in shaping cancer cell behavior. Despite being a proof-of-concept, these results underscore the practical utility of our lineage-informed framework for generating testable mechanistic hypotheses. Given the general applicability of this analytical approach, we expect `MiTo` to provide key biological insights in a wide range of physiological and pathological contexts.

Finally, to contextualize the performance of `MiTo` and MT-scLT in the broader landscape of scLT technologies, we systematically compared single-cell phylogenies across 4 independent scLT platforms. Our analyses revealed that clade support in MT-phylogenies is strongly influenced by tree depth, and the presence of high-confidence MT-SNVs. While MT-SNV can robustly mark distinct cellular clones across a MT-phylogeny, their sparsity and lower character diversity — relative to nuclear SNVs or Cas9-based barcodes — limits robust detection of higher-resolution evolutionary events. Thus, we argue that MT-phylogenies can represent useful scaffolds for clonal inference, but not detailed genealogical histories. While this level of phylogenetic resolution can be successfully exploited in a variety of contexts (e.g., cancer evolution, aging), studies focusing on faster cellular dynamics (e.g., early embryogenesis) should employ higher-resolution approaches – despite limited scalability, multi-omic compatibility, and/or dependency from exogenous cell barcoding. We propose that future scLT systems jointly profiling lineage markers from both mitochondrial and nuclear genomes within the same multi-omic assay could overcome these limitations. This technological development will likely represent the next outstanding challenge for the field.

In conclusion, this work introduces new tools, benchmarking datasets, and practical guidelines for MT-scLT data analysis, validating the its accuracy at unprecedented scales. This study also contributes to the ongoing debate surrounding MT-scLT application range, providing a quantitative reference for scLT system selection, and highlighting key areas of future technological development.

Evolution has long fascinated scientists, providing a unifying theoretical framework that links molecular processes to organismal and population-level phenomena. We anticipate that `MiTo` will substantially advance the study of somatic evolution across a wide range of biological contexts.

## Materials and Methods

This section is divided in: Computational Methods (new tools developed), Data generation (new data generated), and Data Analysis (analysis of newly generated and publicly available data).

### Computational methods

This section gives an overview of the new tools developed within this work: `MiTo and nf-MiTo.`See Supplementary Information – `MiTo` toolkit for details.

### MiTo

The `MiTo` python package introduces novel methods for: i) cell and MT-SNVs filtering; ii) MT-genotyping; iii) Cell-cell distance calculation in MT-SNVs space; iv) Clonal inference from MT-phylogenies. These and other functionalities (e.g., dimensionality reduction, visualization, other utils) are implemented into `scverse-`compatible APIs (see `MiTo’s` <u>documentation</u> for individual function and classes, tutorials, and use cases). Here we describe the main methods developed in `MiTo`. For each method and function cited, default arguments have been set as described in Supplementary Information – `MiTo` benchmark.

### Cell filtering

Implemented in the `mito.pp.filter_cells` function - `MiTo` provides 2 cell filters: i) “filter1”, filtering cells with mean MT-genome (all genome) site coverage >= 20 consensus UMIs; and ii) “filter2”, filtering cells with median (MAESTER-)target site coverage >= 25 consensus UMIs, and fraction of target site covered >= 0.75. For additional information on cell filtering, see Supplementary Information – `MiTo` benchmark – Cell filtering.

Note that, since these cell filters require coverage information across all MT-genome sites, they can be applied only to `MiTo` and `maegatk` AFMs - for which this information is available. See Supplementary Information – `MiTo` benchmark – Raw reads pre-processing and AFM filtering, for details about how this issue has been handled to benchmark different raw MT-data pre-processing pipelines.

### MT-SNVs filtering

MT-SNVs selection is a fundamental step of MT-scLT, as the majority of candidate MT-SNVs detected may be sequencing errors, technical artifacts, or un-informative variants. To retain only “lineage-informative” MT-SNVs, the `mito.pp.filter_afm` function provides surgical MT-SNVs annotation and selection capabilities. Specifically, this function:

1. Annotates MT-SNVs with key summary statistics – previously adopted for MT-SNVs filtering in ^40^ and ^42^
2. Applies a very loose, “baseline” filter - mean site coverage>=5, mean quality>=30, and cells with non-zero AF >=2 – to filter out very unlikely MT-SNV candidates
3. Applies either the main `MiTo` filter - which re-adapt and refine MT-SNV selection criteria from both ^40^ and ^42^ - or other four previously published MT-SNVs filters - `CV`^41^, `miller2022`^40^, `weng2024`^42^, `MQuad`^65^ - to filter putative MT-SNV candidates
4. Filters out common dbSNP variant and RNA-editing events, as in ^55^
5. Calls MT-genotypes – see Materials and Methods – Computational methods – `MiTo` – MT-genotyping - and compute cell-cell distances in the resulting binarized MT-SNV space – see Materials and Methods – Computational methods – `MiTo` – Cell-cell distances in MT-SNV space
6. Filters out MT-SNVs that are not-significantly auto-correlated with cell-cell distances in MT-SNV space (i.e., Moran’s I permutation test pvalue <= 0.05)
7. Filters cells that are positive for at least one of the remaining, high-quality MT-SNV
8. Computes metrics to evaluate properties of the filtered MT-SNV space - see Supplementary Information – `MiTo` toolkit – `nf-MiTo` metrics

`mito.pp.filter_afm` is used to annotate and filter all the MT-SNVs subsets characterized through this work. Note that `samtools` and `freebayes` AFMs - which return already filtered MT-SNV call-sets - were minimally filtered - given the few filtered MT-SNVs, and the lack of MT-SNVs quality and depth statistics - while for `cellsnp-lite` AFMs we used either the `MiTo` filter or the `MQuad` filter, as recommended by the authors. `MiTo` preprocessing and `maegatk` AFMs were filtered with the `MiTo` – or alternatively the `MQuad` - filter, which select variants with:

- Mean variant quality >= 30
- Fraction of negative cells >= 0.2
- Allelic Frequency of confident detection >= 0.02
- Number of +cells >= 5
- Number of confident detection events >= 2
- Mean number of (consensus) UMIs supporting the alternative allele in +cells >= 1.25
- Mean number of (consensus) UMIs on the variant site >= 10

See Supplementary Information – `MiTo` benchmark – Raw reads pre-processing and AFM filtering additional information on default arguments values and alternative filters.

### MT-genotyping

For a given candidate MT-SNV site, all pre-processing pipelines yield alternative and total UMI counts across cells, 𝑎𝑑 and 𝑑𝑝 - 𝑎𝑑, 𝑑𝑝 ∈ ℕ^1*xN*^, with 𝑁 = 𝑛 𝑐𝑒𝑙𝑙𝑠. For a given MT-SNV, the `vanilla` strategy adopted in ^42^ assigns genotype 1 (i.e., MT-SNV presence) to cell 𝑖 if 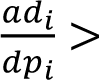 *t_vanilla_* and 𝑎𝑑_*i*_ ≥ 𝑚𝑖𝑛𝐴𝐷, and 0 (i.e., MT-SNVs absence) otherwise. Throughout this work, 𝑡_*vanilla*_ has been always set to 0, while 𝑚𝑖𝑛𝐴𝐷 has been set to either 1 or 2. Conversely, `MiTo` genotyping strategy leverages statistical modelling of 𝑎𝑑 and 𝑑𝑝 counts for accurate estimation of MT-genotypes, effectively taking into account the background noise, in single-cell MT-SNVs data. This is performed by re-adapting the binomial mixture modeling approach introduced by `Mquad` ^65^ to MT-SNVs genotyping, rather than MT-SNVs selection. Specifically, for each MT-SNV, `MiTo` fits the same 2-components mixture model of `MQuad` (implemented in the `MixtureBinomial` class from the `bbmix` package, https://github.com/StatBiomed/BBMix), with the two binomial components representing either the background signal (present also in True Negative cells. Component 0, or equivalently, the 0 genotype) and the true biological signal (ascribed only to True Positive cells. Component 1, or equivalently, the 1 genotype). Then, `MiTo` leverages Bayesian statistics to calculate each cell genotype posterior probabilities, 𝛾_*i*0_ and 𝛾_*i*1_ - see Supplementary Information – `MiTo` toolkit – `MiTo` structure and methods – MT-genotyping for mathematical details. For each MT-SNV, genotype 1 is assigned to cells with 𝛾_*i*1_ > 𝑡_*prob*_ and 𝛾_*i*0_ < 1 − 𝑡_*prob*_, while genotype 0 is assigned to all the other cells. By estimating the background signal associated with the observed alternative UMI counts `MiTo` achieves accurate genotyping, especially for challenging low-detection/high-prevalence MT-SNVs^47^. Since this probabilistic approach performs well only with consistent numbers of detection events (i.e., 𝑎𝑑_*i*_ > 0), the `mito.pp.call_genotypes` function allows flexible tuning of 𝑡_*prob*_, and the minimal cell prevalence required for a MT-SNVs variant to be genotyped this way (i.e., very low-prevalence variants are still genotyped with the simpler `vanilla` method, to avoid too sparse data fitting problems). See Supplementary Information – `MiTo` benchmark – MT-genotyping for 𝑡_*prob*_ and 𝑚𝑖𝑛_𝑐𝑒𝑙𝑙_𝑝𝑟𝑒𝑣𝑎𝑙𝑒𝑛𝑐𝑒 default values choice (0.7 and 0.05, respectively).

### Cell-cell distances in MT-SNV space

The `mito.pp.compute_distances` function accept several distance metrics (see `MiTo` APIs documentation). Across this study we used either an unweighted or weighted jaccard distance, calculating each MT-SNV weight as its median AF in +cells. This weighting strategy makes the jaccard distance (a strict measure of mutational set overlap between cells) dependent on the strength of MT-SNVs signal, down-weighting too sparsely or too weakly observed variants. Cell-cell distances are also rescaled into the [0,1] interval, to maximize signal to-noise ratio. See Supplementary Information – `MiTo` benchmark – Distances in MT-SNV space for distance metrics benchmarking.

### Clonal inference from MT-phylogenies

`MiTo` uses MT-phylogenies to infer discrete cell clones (i.e., group of cells genetically defined by one or more MT-SNVs, clustering together on the phylogeny, under a least common ancestor). This task is performed into five sequential steps. First, a MT-phylogeny is inferred from a pre-processed AFM (i.e., `mito.tl.build_tree`, a wrapper around `Cassiopeia`^66^ APIs for lineage inference. The default algorithm for tree reconstruction is the distance-based Unweighted Pair-Group Method with Arithmetic Means method, or UPMGA). For each MT-SNV and internal node, the Fisher’s Exact Test is used to calculate an enrichment score (i.e., −𝑙𝑜𝑔_10_𝑝𝑣𝑎𝑙𝑢𝑒), quantifying how much is the variant has been exclusively detected in a given node’s clade. Then, a recursive function inspired by ^16^ is used to traverse the tree in a depth-first manner, to identify clades that: i) have at least one MT-SNV with enrichment score > *mut_enrichment_treshold*, and ii) for which the the median cell-cell similarity is < *similarity_treshold.* At each recursion, if such a clade is found, the corresponding MT-SNVs is removed from the set of available MT-SNVs, and another recursion it attempted to check whether both children clades clades satisfy i) and ii), otherwise all cells from the original clades are collected as a mitochondrial clone (MT-clone, with the corresponding MT-SNVs assigned). Once all of such clades have been collected, a redundant (i.e., similarity threshold between clones’ genotypes < *merging_treshold*) clones are merged, starting from the tiniest ones. The procedure stops when no more redundant clones are found. Finally, some diagnostic checks are performed to check the integrity of MT-clones (i.e., no disjoint set of cells across the phylogeny), and to mark as unassigned (i.e., NA) cells that were not assigned to any MT-supported clade.

The clonal inference algorithm described above depends on 3 hyper-parameters (i.e., *mut_enrichment_treshold, similarity_treshold, and merging_treshold*). To achive optimal results, a Grid Search hyper-parameter tuning is performed, leading to different clonal inference solutions, and a heuristic is employed to choose the best performing clonal inference solution, maximizing the average MT-clone silhouette score and cell-cell similarity while minimizing the number of MT-clones. Clonal labels are added to the tree cell metadata, together with additional metrics (e.g., each internal fitness score).

These clonal inference and annotation functionalities are implemented in the `mito.tl.MiToTreeAnnotator` class and its methods. See Supplementary Information – `MiTo` benchmark – Clonal inference to know about alternative methods details, and their benchmark in terms of run-time and memory efficiency.

### nf-MiTo

The `nf-MiTo` pipeline implements `MiTo` methods - together with other, previously published ones - into a flexible, end-to-end data analysis tool. The MT-scLT analysis may start from raw sequencing MT-data – aligned or unaligned MT-reads from the MAESTER protocol^40^ - or from an already preprocessed AFM. A complete description of `nf-MiTo` structure, entry-points, hyper-parameters, and metrics can be found in Supplementary Information – `MiTo` toolkit. See <u>nf-MiTo</u> for usage examples.

### nf-lenti

GBC libraries (both bulk and single-cell reads) were processed with the `nf-lenti` pipeline, which is under preparation in a companion paper. All details will be available at publication date.

### Data generation

This section gives an overview of the scLT experiments and protocols adopted to generate `MiTo` and `nf-MiTo` benchmarking datasets. See Supplementary Information – Data generation for details.

### Dual scLT experiments

We generated 3 new scLT datasets from 3 high-quality biological specimens, all derived from the MDAMB231 Breast Cancer cell line.

#### Plasmids

The Perturb-seq^52^ GBC library (pBA57117,18) was purchased from Addgene and used for both lineage tracing experiments. This vector contains a random 18-nt guide barcode (GBC) between the blue fluorescent protein (TagBFP) and polyadenylation signal sequences. This vector contains puromycin and ampicillin resistance genes and the reporter gene TagBFP constitutively expressed under the control of EF1a promoter. The pLenti CMV Puro LUC (w168–1) was purchased from Addgene. This vector also contains puromycin and ampicillin resistance genes.

#### Cell lines

All cells were cultured in 5% CO2 incubator at 37° C. HEK293T and the metastatic human TNBC cell line MDA-MB-231 were purchased from the ATCC and cultured in DMEM (EuroClone), supplemented with 10% South American FBS, 2 mmol/L L-glutamine, and 100 U/mL penicillin–streptomycin. All the cell lines were tested for Mycoplasma contamination routinely. All the cell lines were split once they reached approximately 80% confluence and cultured in vitro for no more than 10 passages after thawing. Puromycin (2 µg/mL, for 72 hours) was administered for GBC+ cells selection. Transient transfection with Lipofectamine TM was performed uniquely to transfect the Perturb-seq GBC library in HEK293T.

#### Mice

Female NOD/SCID Il2-Rg null (NSG) mice were purchased from Charles River Laboratory and housed under pathogen-free conditions at 22° C±2° C, 55%±10% relative humidity, and with 12 hours d/light cycles in mouse facilities at the European Institute of Oncology–Italian Foundation for Cancer Research Institute of Molecular Oncology (Milan, Italy) campus. In vivo studies were performed after approval from our fully authorized animal facility and our institutional welfare committee and notification of the experiments to the Ministry of Health (as required by the Italian Law (D.L.vo 26/14 and following amendments); IACUC numbers: 833/2018, 679/2020), in accordance with EU directive 2010/63.

#### *In vitro* scLT experiment (MDA_clones dataset)

Single-cells were isolated into 96-well plates by limiting dilution and expanded for ∼30 days. The resulting colonies were infected with unique barcodes (n=8 distinct colonies were infected with 8 unique lentiviral species). After selection, barcoded clones were mixed at known ratios, FACS-sorted for tagBFP expression, subjected to library preparation and sequenced.

#### *In vivo* scLT experiment (MDA_PT and MDA_lung samples)

MDA-MB-231 cells (3 x 10^0^ per plate) were infected with the Perturb-seq GBC library at MOI = 0.3 and selected as described above. For each mouse, 200,000 MDA-MB-231 cells were resuspended 1:1 in 30-µL PBS and growth-factor reduced Matrigel, and injected in the ninth mammary gland of female NSG mice. PTs were monitored by calliper 3 times a week and PT collection was performed ∼30 days after inoculation. Metastatic infiltration was monitored by IVIS-Lumina. Lung metastases were harvested at mice sacrifice, ∼30 days post PT resection. For this work, we selected a PT-lung couple from the scLT experiment described above (involving many other mice and treatment conditions), and processed as described above.

### Library preparation and sequencing

#### 10x cDNA and gene expression (GEX) library preparation and sequencing

FACS-sorted cells were counted via 1:1 mixing with erythrosin B, then pelleted at 2000 rpm for 5 minutes at 4 degrees, resuspended in to reach ∼1000 cells/µL, and counted again. Cell suspensions (∼5000-6000 cells per sample) were loaded into the 10x Chromium chip, following the <u>10x 3’ v3</u> protocol for cDNA production and gene expression (GEX) library preparation. Resulting full length cDNA and GEX libraries were quality-controlled through BioAnalyzer (Agilent). GEX libraries were sequenced with the NovaSeq 6000 Sequencing System (∼50k reads per cell, ∼300M reads per sample). The final layout of GEX (paired-end) reads is:

- R1 (28 bp): CB (16bp) + UMI (12bp)
- R2 (91 bp): endogenous transcript amplified region (3’)

#### Perturb-seq (GBC) bulk library preparation and sequencing

Bulk sequencing of DNA-integrated GBC was performed to retrieve a comprehensive whitelist of (error-corrected) lentiviral barcodes for MDA_PT and MDA_lung samples. Specifically, 200 ng of genomic DNA from approximately 10^0^ cells per sample (mixed with spike-in controls) were amplified and sequenced as in ^67^.

#### Perturb-seq (GBC) library preparation and sequencing

20 ng of each GEX library was used to generate the corresponding GBC library. See supplementary methods and ^67^ for the detailed procedure, primer sequences and PCR cycles. The final layout of GBC (paired-end) reads is:

- R1 (28 bp): CB (16bp) + UMI (12bp)
- R2 (91 bp): Perturb-seq transcript amplified region

#### MAESTER (MT) library preparation and sequencing

The mitochondrial transcripts (MT-) library was generated from 10x barcoded cDNA following the original MAESTER ^40^ protocol. As for the GBC library, a semi-nested PCR approach is adopted. See Supplementary Information – Data generation – MAESTER library preparation and sequencing for the PCR protocol and cycles.

Final MAESTER libraries were quality-controlled by BioAnalyzer, and sequenced with the NovaSeq 6000 Sequencing System (∼50k reads per cell, ∼300M reads per sample).

The final layout of MT-(paired-end) reads is:

- R1 (28 bp): CB (16bp) + UMI (12bp)
- R2 (164 bp): MT-transcript amplified region

### Data analysis

This section describes the computational analyses presented in this work. See Supplementary Information – `MiTo` benchmark and Supplementary Information – Multi-omic analysis of longitudinal breast cancer clones for details.

### Raw data pre-processing

For all newly generated dataset, GEX, MT and GBC libraries - paired-end fastq files, with R1 and R2 reads in the standard 10x genomic format, see Materials and Methods – Data Generation – Library preparation and sequencing - were pre-processed with `nf-MiTo` (GEX, MT) and `nf-lenti` (GBC) pipelines – see Materials and Methods – Computational methods. Since local tuning of `nf-MiTo` hyper-parameters required “raw”, unfiltered Allele Frequency Matrices (AFMs) from different pre-processing pipelines – see Supplementary Information – `MiTo` benchmark – Raw reads pre-processing and AFM filtering – our data were pre-processed as follows:

1. GEX and MT data were pre-processed with `nf-MiTo PREPROCESS` - enabling the `– -raw_input_data_type` = “fastq” option, see Supplementary Information – `MiTo` toolkit – `nf-MiTo` entrypoints. This first run yielded `maegatk`^40^ Allele Frequency Matrices (AFMs) and GEX matrices filtered for quality-controlled cell barcodes (CBs) - see Materials and Methods – Data analysis – Cell QC - together with other intermediate outputs - e.g., quality-controlled CBs, aligned MT-reads.
2. Quality-controlled CBs were used to pre-process GBC libraries with `nf-lenti` - default configuration. Importantly, to enhance the robustness of `nf-lenti` clonal assignment, reference GBC whitelists were provided for each sample. This included the known list of GBC sequences for MDA_clones, and the two lists of GBC sequences detected in bulk, targeted-DNA sequencing for MDA_PT and MDA_lung samples – see Materials and Methods – Data Generation – Library preparation and sequencing. `nf-lenti` yielded robust GBC clonal labels that were added to the `maegatk` AFMs and GEX matrices cell metadata obtained previously – note, only cells robustly assigned to a single GBC-clone were filtered. Unassigned cells or cells assigned to multiple GBC-clones were filtered out.
3. The remaining AFMs - `cellsnp-lite`^68^`, MiTo, freebayes`^69^`, samtools`^70^ pipelines – were obtained by pre-processing aligned MT-reads, quality-controlled CBs, and GBC-containing cell metadata. Again, this task was performed with `nf-MiTo PREPROCESS`, enabling the `–-raw_input_data_type` = “mitobam” option and the

`–-pp_method`, `--path_meta` hyper-parameters – see Supplementary Information

– `MiTo` toolkit – `nf-MiTo` hyper-parameters.

These pre-processing operations yielded “raw”, unfiltered AFMs from 5 MT-reads pre-processing pipelines, each including only GBC-labelled and (GEX) quality-controlled cells. This level of pre-processing corresponds to `nf-MiTo` step i (Fig. 1), with the addition of GBC labels in AFM cells metadata. Fig. 2a-c show samples summary statistics - number of cells, number GBC clones, GBC-clonal complexity, mean MT-genome coverage - considering `maegatk` AFMs, the best performing pre-processing pipeline in our benchmark – see Supplementary Information – `MiTo` benchmark – Raw reads pre-processing and AFM filtering. Fig. 2-5 *always* show lower numbers of cells and GBC-clones, due to the additional cell and MT-SNVs filtering operations – `nf-MiTo` step ii - needed for robust lineage inference – see Supplementary Information – `MiTo` benchmark.

### Cell QC

`nf-MiTo` implement a simple cell Quality Control to exclude low-quality cells – according to their gene expression profile. Specifically, good quality cells must meet the following criteria:

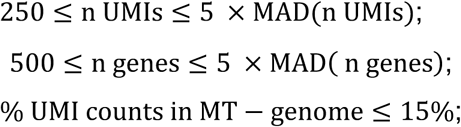

with MAD: Median Absolute Deviation.

### nf-MiTo tuning

`nf-MiTo` computes n=19 diagnostic metrics, and allow fine control of the whole scLT workflow with its n=53 command-line parameters - see Supplementary Information – `MiTo` toolkit - `nf-MiTo` metrics, and Supplementary Information – `MiTo` toolkit - `nf-MiTo` hyper-parameters. After development of the main `MiTo` algorithms, we wanted to find the best pipeline approach for MT-scLT data analysis - the best combination of hyper-parameters for each `nf-MiTo` step. To achieve this, we employed a local tuning strategy. First, a single default configuration was set for all step, considering preliminary lineage inference performance evaluations – see Supplementary Information – `MiTo` benchmark – Preliminary assessments. Then, for each sequential step, we perturbed the most important hyper-parameters and quantified gains and losses in performance, leaving all the other hyper-parameters unchanged. After multiple rounds of local tuning and hyper-parameters updates, the best hyper-parameters were finally set as `nf-MiTo` default configuration. See Supplementary Information – `MiTo` benchmark for details.

### Clonal inference benchmark

`nf-MiTo` tuning demonstrated that lineage inference (i.e, step v, the step at which lineage inference performance is actually evaluated) is deeply influenced by other pre-processing (step i-iv) steps, and their associated options. Therefore, we assessed lineage inference only on AFMs with reasonable phylogenetic signal. In practice, for each sample we selected only the best (n=5) AFMs – i.e., filtered AFMs with the highest concordance between inferred and ground truth clones (as measured by ARI) and decent cellular yield. See Supplementary Information – `MiTo` benchmark – Clonal inference for sample-specific thresholds – and provided them to 4 alternative clonal inference methods: i) `MiTo`; ii) `leiden`, iii) `vireoSNP`; iv) `CClone.` This choice was made to avoid un-necessary (and un-informative) high numbers of benchmarking trials, and to focus on each method optimization, rather than other up-stream data processing steps. See Supplementary Information – `MiTo` benchmark – Clonal inference for methods’ specifics and benchmarking details.

### Multi-omic analysis of longitudinal Breast Cancer clones

#### Cell state annotation

MDA_PT and MDA_lung (PT and lung) gene expression matrices (preprocessed with `nf-MiTo` and `nf-lenti` as described in *GEX and GBC raw data pre-processing*) were quality controlled (as described in *Cell QC*) and merged together. We used `scanpy`^71^ for GEX analysis. First, we assessed the presence of additional low-quality cell clusters. After library size normalization, the top (n=2000) Hyper-Variable Genes (HVGs) were selected for dimensionality reduction (PCA, n components=50), and kNN graph construction (top 30 PCs, k=15). The `leiden` algorithm (resolution 0.1-2) was used cluster cells. Here, two cell clusters with low number of UMIs and detected genes were removed. Then, remaining cells were subjected to another round of pre-processing and clustering. After library size normalization and HVGs selection, we removed cell-cycle correlated HVGs (Pearson’s correlation with MKI67 gene > 0.5), reduced data dimension with PCA (n components=50), and selected the top 20 PCs for kNN graph construction. This graph was used to compute diffusion maps (n components=10), and a denoised kNN graph (k=50). This final kNN graph was used for visualization (`UMAP`^72^ algorithm, Fig. 4c), and clustering (resolution 0.05-0.15). Differential Expression analysis (DE, Wilcoxon’s rank test) and Gene Set Enrichment Analysis^73^ (GSEA, implemented in `gseapy`^74^) were used to annotate unsupervised clusters into broad cell states.

#### Lineage inference

Pre-processed AFMs from PT and lung – see Materials and Methods – Data analysis – Raw data pre-processing - were merged and filtered for cells with annotated cell states – see Materials and Methods – Data analysis – PT-lung multi-omic analysis – Cell state annotation. This merged AFM was used as input for lineage inference with `nf-MiTo` – the `INFER` entrypoint with default setting was used, see Supplementary Information – `MiTo` toolkit – `nf-MiTo` entrypoints for details. The resulting MT-phylogeny and -clones (Fig. 4b) were used for all downstream analyses in Fig 4. and Supp. Fig. 16-17.

#### Clonal bias and cell state transition dynamics

For each each clone (MT- or GBC-), cell state bias was visualized in term of cell frequencies (i.e., Fig. 4d, the number of cells populating each cell state normalized by the total number of cells) and enrichment scores (−𝑙𝑜𝑔_10_𝑝𝑣𝑎𝑙𝑢𝑒 from right-tailed Fisher’s exact test). Cell state transitions between PT and lung were computed with `moslin`^56^. `moslin` solves a Fused Gromov-Wassertein optimal transport problem to infer a transition map that represent the (average) probability for a cell in cell state 𝑖 at timepoint 𝑡_*i*_ to reach cell state 𝑗 at timepoint 𝑡_*i*_. `moslin` is implemented in a broad toolkit for single-cell optimal-transport, `moscot`^75^ (i.e., `moscot.probles.time.LineageProblem` class). For this comparative analysis, we inferred the GBC- and MT-transition map independently, using different combinations of cell-cell distances in gene expression and lineage space. Specifically, we used euclidean distances in (PC) gene expression space, combined with either MT- (weighted jaccard distances in MT-SNV space) or GBC- (jaccard distance in binarized GBC space) lineage distances. Transition map inference and aggregation (Fig. 4e) was performed with default parameters, as indicated in `moslin` tutorials.

#### Transcriptional determinants of clonal fitness

`mito.tl.nb_regression` implements the Negative Binomial (NB) regression framework introduced in ^42^. Specifically, a generalized linear model is employed to model the raw, pseudo-bulk counts of gene 𝑦 as a function of other covariates. First, for each clone (min n cells=30) we created pseudo-bulk samples, partitioning its cells into one (or more) cell samples of the same size (n cells=30), and aggregating raw UMI counts by sum (`mito.tl.agg_pseudobulk`). Then, for each gene, we used these pseudo-bulk data to fit two NB negative binomial models, including either (z-scored) clone size or clone branchedness^31^ as predictor variables. For each fitted model, the estimated slope and pvalue (i.e., the strength and robustness of each gene-clone feature statistical association, respectively) were used to rank individual genes (Fig 4g) and as input to GSEA, for further biological interpretation.

#### Memory and plasticity of gene expression programs

Raw GEX data were used to infer gene regulatory networks (or regulons) with `pySCENIC (`default parameters). `scanpy.tl.score_genes` was used to compute each cell regulon activity score, which were then rescaled (z-scored, Fig 4m). The Moran’s I statistic was used to quantify the level of spatial autocorrelation of each regulon with MT-distances (Fig 4m-n, weighted jaccard distances between MT-genotypes, computed by `nf-MiTo`).

### scLT systems comparative analysis

#### Public datasets

RedeeM data were downloaded and pre-processed as described in https://github.com/caleblareau/redeem-reanalysis-reproducibility. Pre-processed scWGS data were downloaded from https://github.com/margaretefabre/Clonal_dynamics. Cas9 data were downloaded and pre-processed as described in https://www.sc-best-practices.org/trajectories/lineage_tracing.html.

For all re-analyzed samples, custom scripts leveraging the `mito.io.make_AFM` function were employed to assemble properly formatted AFMs - from externally pre-processed character matrices (and priors, for Cas9 data).

#### Linage inference

Properly formatted AFMs – see Materials and Methods – Data Analysis – scLT comparative analysis – Public datasets - were provided to `nf-MiTo INFER` entrypoint for lineage inference - see Supplementary Information – `MiTo` toolkit - `nf-MiTo` entry-points. The resulting phylogenies (Fig. 5b) and tree metrics were used for downstream comparison (Fig 5c). Originally pre-processed character matrices (i.e., filtered cells and characters) and priors (Cas9 datasets) were used for all published datasets. To avoid potential biases arising from different tree building algorithms, the general-purpose, distance-based UPMGA algorithm was used to infer all single-cell phylogenies. For scWGS data, all colonies and SNVs were retained for phylogeny inference and cell genotypes provided by the authors were used to compute cell-cell distances (unweighted jaccard distance). The same holds for Cas9 data (i.e., no additional cell and character filtering, after pre-processing suggested by the authors^31^). Cell-cell distances were estimated with a weighted hamming distance, developed in ^66^ for Cas9 INDELs characters. For MAESTER data, the default `MiTo` workflow was used on `maegatk` AFMs (i.e., `MiTo` AFM filtering, genotyping and weighted-jaccard distance computation), as described in Materials and Methods - Computational Methods – `MiTo`. For RedeeM data, character matrices pre-processed from the authors underwent additional filtering, given recent concerns about the quality of original RedeeM data, and the additional benefits demonstrated by `MiTo` cell and MT-SNVs filtering on MAESTER data. In practice, we first filtered only cells with mean coverage (endogenous UMIs) >=20 across the entire MT-genome (“filter_1” in `MiTo` cell filtering), and applied the following shallow filters (`MiTo` baseline MT-SNV filter) to the original RedeeM^42^ MT-SNVs:

- Mean site coverage (DP) >= 5
- n +cells >= 2
- No MT-SNVs with more than one alternative allele

In addition, we discarded all MT-SNVs flagged as common variants/RNA edits in REDdb and dbSNP databases (see `MiTo` MT-SNVs filters). Then, binary genotypes were assigned to all remaining cells and MT-SNVs (`--bin_method` = `vanilla`, `–-t`_`vanilla` > 0 and `-- min_AD` = 1, see Supplementary Information – `nf-MiTo` hyper-parameters). Only cells positive for at least one of these filtered MT-SNVs were retained for phylogeny inference. Since all of these filtering steps should discard putative non-somatic MT-SNVs from character matrices, we did not used the empirical priors of the original RedeeM study to correct cell-cell jaccard distances and downweight frequently occurring MT-SNVs – i.e., we used standard jaccard distance.

### Data availability

Newly generated data will be made public at publication date.

### Code availability

`MiTo`: https://github.com/andrecossa5/MiTo

`nf-MiTo`: https://github.com/andrecossa5/nf-MiTo

Reproducibility code: https://github.com/andrecossa5/MiTo_benchmark_repro

