## Supplementary Information for "MiTo: tracing the phenotypic evolution of somatic cell lineages via mitochondrial single-cell multi-omics"

This section provides technical details for computational methods, analyses and results described in this work, divided into 5 main sections: i) `MiTo` toolkit; ii) Data generation; iii) `MiTo` benchmark; iv) Multi-omic analysis of longitudinal Breast Cancer clones;

#### `MiTo` toolkit

Here we provide detailed information about the `MiTo` python package – code organization and methods - and the `nf-MiTo` Nextflow pipeline – structure, entry-points, hyper-parameters and evaluation metrics (Fig.1).

##### `MiTo`

`MiTo` (v.0.0.2) is structured as a typical `scverse` package designed for user-friendly and interactive use. `MiTo` functions and feature-rich APIs can be imported from 4 main collections of modules: `.io`, functions to create AFMs, convert data format, read and write operations; `.pp`, preprocessing utils; `.pl`, plotting utils; `.tl`, tools for tree building and lineage-phenotype integrative analyses; `.ut`, general purpose utils. This standardized organization ensures easy access to each module, function and class for end-users of different computational expertise. Importantly, the package includes extensive tutorials and pre-processed dataset, allowing users to quickly and efficiently learn `MiTo` functionalities for real-world data applications. `MiTo` is open-source, and distributed by pypi.

MT-genotyping derivation

This section describes the statistical foundation of `MiTo` genotyping (for a general overview, see Materials and Methods - Computational Methods – `MiTo` - MT-genotyping).

Assuming diploid status across all cells in a population, the genotype of a nuclear SNVs in a given cell cells can be either homozygous WT (00 state, AF of the mutated allele=0), homozygous MUT (11 state, AF ot the mutated allele=1), or heterozygous (01 state, AF ot the mutated allele=0.5). On the contrary, MT-SNVs display a nearly continuous spectrum of allelic frequencies, that can be binarized with two genotype states: 0 → absence of a MT-SNVs, 1→ presence of the MT-SNVs. It has been previously shown that discrete metrics are more robust to scRNA-seq technical noise than continuous ones with regards to quantifying pairwise cell-cell genetic (dis-)similarity. However, these former metrics need accurate MT-SNVs genotyping (i.e., AFM binarization). In this work, we implemented two binarization methods: vanilla and `MiTo`. For a given MT-SNV site, one can retrieve alternative and total UMI counts observed across the cell population, $ad$ and $dp$ ($ad$ , $dp \in \mathbb{N}^{1xN}$, with $N = n\ cells$). For a given MT-SNV, the `vanilla` method assigns genotype 1 to cell $i$ if $ad_i \ / \ dp_i > t_{vanilla}$

and $ad_i >= minAD$. Throughout this work: $t_{vanilla}$ has been set to 0, while $minAD$ has been set to either 1 or 2. On the other hand, `MiTo` genotyping leverages the statistical modelling of $ad$ and $dp$ counts to assign binary genotypes to each cell. In particular, we re-adapted the probabilistic approach introduced by `Mquad` to MT-SNVs genotyping. To do this, we assumed that, $ad$ , $dp$ counts are generated by weighted sampling of two binomial distributions, representing the background (component 0) and the true positive signal (component 1), respectively. Under this assumption, denoting with $ad_i$ and $dp_i$ the alternative and total UMI counts for cell *i* respectively, the probability $P$ of observing exactly $ad_i$ alternative UMI counts for cell $i$ is defined as:

$$P: \mathbb{N}^+ \rightarrow [0,1]; \;\; ad_i \rightarrow f(ad_i \mid dp_i, \Theta)$$

$$P(ad_i \mid dp_i, \Theta) := \sum_{k=0}^{K} \pi_k \cdot Binomial(ad_i \mid dp_i, p_k) \quad \text{Eq. 1}$$

with:

$$Binomial(x \mid n, p) := \binom{n}{x} p^x (1-p)^{n-x}$$

$K = 1$, with $k = 0$ background component and $k = 1$ true positive signal component

$\Theta = [p_k, \pi_k] \in \mathbb{R}^{2(K+1)}$, parameter vector of the model

$p_k \in [0,1]$, success rate for component *k*

$\pi_k \in [0,1]$, mixing weight for component *k*

$$\sum_{k=0}^{1} \pi_k = 1$$

This probabilistic model can be fitted to the observed values of the $ad$ , $dp$ UMI counts by maximizing the total likelihood $L(\Theta \mid ad, dp)$:

$$L : \mathbb{R}^{2(K+1)} \rightarrow \mathbb{R}^+;$$

$$L := \prod_{i=1}^{N} P(ad_i \mid dp_i, \Theta) \qquad \text{Eq. 2}$$

By taking the logarithm of both sides and unfolding the definition of the binomial distribution probability mass function, Eq. 2 becomes:

$$logL = log(\prod_{i=1}^{N} P(ad_i \mid dp_i, \Theta))$$

$$= \sum_{i=1}^{N} ( log \binom{ad_i}{dp_i} + log(\sum_{k=0}^{K} \pi_k \cdot p_k^{ad_i} \cdot (1 - p_k)^{dp_i - ad_i}) ) \quad \text{Eq. 3}$$

Maximization of this $logL$ - equivalent to likelihood maximization given the monotonicity of the logarithm function - can be nicely achieved via Expectation-Maximization (EM), a Bayesian optimization approach that is capable of handling hidden variables through Expectation (E) and Minimization (M) steps. First, at each E-step, Bayes Theorem is used to compute the posterior probabilities of latent (*hidden*) variables $Z_i \in \mathbb{N}^{1xN}$, i.e., the cell memberships to either background ($k = 0, Z_i = 0$) or true positive signal ($k = 1, Z_i = 1$) components (i.e., cell genotypes). Given the definition of posterior probability of A given B, $P(A \mid B)$, from Bayes Theorem:

$$P(A \mid B) := \frac{P(B \mid A)\, P(A)}{P(B)}$$

Considering:

$$A = P(Z_i = k)$$
$$B = [ad_i, dp_i, \Theta^\star]$$
$$\Theta^\star \text{ current estimate for } \Theta$$

the posterior probability of cell $i$ assignment to the k-th component is defined as:

$$\gamma_{ik} := P(Z_i = k \mid ad_i, dp_i, \Theta^\star) = \frac{P(\,ad_i \mid dp_i, Z_i = k, \Theta^\star)\, P(\,Z_i = k \mid \Theta^\star)}{P(\,ad_i \mid dp_i\,, \Theta^\star)} \quad \text{Eq. 4}$$

With:

$$P(Z_i = k \mid \Theta^\star) = \pi_k^\star$$
$$P(ad_i \mid dp_i, Z_i = k, \Theta^\star) = Binomial(ad_i \mid dp_i, p_k^\star)$$
$$P(\,ad_i \mid dp_i, \Theta^\star) = \sum_{k=0}^{K} \pi_k \cdot Binomial(ad_i, dp_i, p_k^\star), \text{ from Eq. 1}$$

Eq. 4 becomes:

$$\gamma_{ik} = \frac{\pi_k^\star \cdot Binomial(\,ad_i \mid dp_i, p_k^\star)}{\sum_{k=0}^{1} \pi_k \cdot Binomial(ad_i, dp_i, p_k^\star)} = \frac{\pi_k^\star \cdot p_k^{ad_i} \cdot (1-p_k^\star)^{dp_i - ad_i}}{\sum_{k=0}^{K} \pi_k^\star \cdot p_k^{ad_i} \cdot (1-p_k^\star)^{dp_i - ad_i}} \quad \text{Eq.5}$$

These estimated $\gamma_{ik}$ are then used to update $\Theta^{\star}$ in the following M-step. Specifically, Eq. 3 describes the $logL$ of the mixture model, *without* including unknown cell membership, explicitly. If these memberships were known (omitting terms not dependent on model parameters $\Theta$) Eq. 3 would simplify from:

$$logL(\Theta)_{complete} = \sum_{i=1}^{N} log(\sum_{k=0}^{K} \pi_k \cdot p_k^{ad_i} \cdot (1 - p_k)^{dp_i - ad_i})$$

to:

$$logL(\Theta)_{complete} = \sum_{i=1}^{N} \sum_{k=0}^{K} \delta_{ik} \, (log\pi_k + ad_i logp_k + (dp_i - ad_i)(1 - p_k)) \quad \text{Eq. 6}$$

where $\delta_{ik}$ is the Kroenecker delta defined as:

$$\delta_{ik} = 1 \text{ if } Z_i = k, \text{ and 0 otherwise}$$

While $Z_i$ (and therefore $\delta_{ik}$) are unknown, the expected values $\gamma_{ik} := E[\, Z_i = k \mid ad_i, dp_i, \Theta^{\star}]$ are estimated in the E-step. Thus, substitution of $\delta_{ik}$ with $\gamma_{ik}$ in Eq. 6, gives:

$$logL(\Theta)_{complete} = \sum_{i=1}^{N} \sum_{k=0}^{K} \gamma_{ik} \, (log\pi_k + ad_i logp_k + (dp_i - ad_i)(1 - p_k)) \quad \text{Eq. 7}$$

$logL(\Theta)_{complete}$ is not a sum over all $N$ observations and $K$ model components without involving log-sum terms and with nicely separated model parameters. Thus, analytic methods for direct maximization can be readily used to update $\pi_k$ and $p_k$ separately. New $\pi_k$ values, $\pi_k^{new}$, are computed by solving:

$$argmax \; logL(\pi_k)_{complete} = argmax \; \sum_{i=1}^{N} \sum_{k=0}^{K} \gamma_{ik} log\pi_k + const \quad \text{Eq.8}$$
$$subjected \; to: \sum_{k=0}^{1} \pi_k = 1$$

which gives:

$$\pi_k^{new} = \frac{1}{N} \sum_{i=1}^{N} \gamma_{ik} \text{ , with } k \in \{0,1\} \qquad \text{Eq.9}$$

Instead, new $p_k$ values, $p_k^{new}$, are computed by solving:

$$argmax\ logL(p_k)_{complete}\ =$$
$$= argmax\ \sum_{i=1}^{N}\sum_{k=0}^{K}\gamma_{ik}(ad_i logp_k + (dp_i - ad_i)(1-p_k)) + const \quad \text{Eq.10}$$

which gives:

$$p_k^{new} = \frac{\sum_{i=1}^{N}\gamma_{ik}\cdot ad_i}{\sum_{i=1}^{N}\gamma_{ik}\cdot dp_i}, \text{ with } k \in [0,1] \quad \text{Eq.11}$$

Crucially, the EM steps alternate between estimating posterior probabilities for genotypes $Z$ and updating model parameters until convergence of log-likelihood values is reached. Across this optimization path, $Z$ plays the fundamental role of bridging the gap between the observed data and the structure of the probabilistic model. However, since `MQuad` only uses the likelihood of this mixture model to rank individual MT-SNVs, the cell genotypes $Z$ and their *expected* values $\gamma_{ik}$ are "lost" in the fitting process. To genotype individual MT-SNV, the `MiTo` *genotyping* method fits the same mixture model employed by `MQuad` (implemented in the `MixtureBinomial` class from the `bbmix` package, https://github.com/StatBiomed/BBMix), but then, for each cell, `MiTo` calculates the $Z_{ik} = 0$ and $Z_{ik} = 1$ genotypes posterior probabilities, $\gamma_{i0}$ and $\gamma_{i1}$ . For each MT-SNV, genotype 1 is assigned to cells with $\gamma_{i1} > t_{prob}$ and $\gamma_{i0} < 1 - t_{prob}$, while genotype 0 is assigned to all the other cells. By estimating the background signal associated with the observed alternative UMI counts `MiTo` achieves accurate genotyping, especially for challenging low-detection/high-prevalence MT-SNVs. Since the binomial mixture model performs best with relatively high numbers of detection events, the `mito.pp.call_genotypes` function allows flexible tuning of $t_{prob}$ and the minimal cell prevalence required for a MT-SNVs variant to be genotyped this way. All the other low-prevalence variants are genotyped with the simpler `vanilla` method.

**`nf-MiTo` entrypoints**

`nf-MiTo` implements `MiTo` core functionalities into Nextflow DSL2 modules. This design allows platform-agnostic, portable, and scalable execution of the data processing steps needed for MT-scLT. These features enable efficient and automatic processing of MT-data by end users, and facilitate benchmarking of novel methods by developers (for a general overview, see Materials and Methods - Computational Methods - `nf-MiTo`). Importantly, the modular DSL2 syntax guarantees atomic development and reuse of individual data processing components, a feature that will consent easy integration of novel functionalities in the future. As of today, `nf-MiTo` - default entry-point - is an end-to-end pipeline, with raw sequencing data as input and annotated (MT-)phylogenies as output. However, to achieve more flexibility in real-world scenarios, nf-MiTo provides 5 additional entry-points: i) PREPROCESS; ii) TUNE; iii) EXPLORE; iv) INFER; v) BENCH.

PREPROCESS

The PREPROCESS entry-point takes as input MT-reads from the MAESTER protocol - see Materials and Methods - Data generation - Library preparation and sequencing - MAESTER (MT) library preparation and sequencing; and Supplementary Information - Data generation - and streamlines all the operations needed to process them into an Allele Frequency Matrix (AFM) – a cell x variant matrix, storing raw Allelic Frequencies for all detectable MT-SNVs in a cell subset of interest (Fig.1, nf-MiTo step i).

The --raw_data_input_type hyper-parameter - see Supplementary Information – MiTo toolkit – nf-MiTo hyper-parameters - specifies for the "raw" sequencing data type provided by the user. This can include: i) unaligned GEX- and MT-libraries in .fastq format - "fastq" option -; ii) unaligned MT-libraries in .fastq format, and a list of 10x cellular barcodes (CBs) of interest - "fastq, MAESTER" option - ; and iii) aligned MT-reads in .bam format, and a list of CBs of interest - "mitobam" option. Depending on the raw sequence input, GEX and/or MT-reads are mapped to the reference genome with STAR Solo[1]. When the "fastq" option is enabled, GEX data pre-processing – reads alignment, UMI counting, and cell Quality Control - internally provide the CB list used to filter aligned MT-reads before further processing.

Regardless of the raw sequence input, aligned and filtered MT-reads are always processed into AFMs with one of the 5 alternative pre-processing methods supported by nf-MiTo: maegatk (default method), MiTo, cellsnp-lite, samtools, freebayes. See Supplementary Information - MiTo benchmark - Raw reads pre-processing and AFM filtering for individual methods details, and their benchmark.

In this work, all GEX and MT-data were processed with nf-MiTo PREPROCESS. To do this, the hg38 reference genome and its transcripts annotations were downloaded from the Cell Ranger Downloads page. Then, this reference genome was complemented with a custom DNA sequence – two constant regions of the transcribed Perturb-seq vector flanking an 18-bp stretch of Ns, mimicking the lentiviral genetic barcode (GBC) - to allow GBC-containing reads alignment and retrieval (i.e., nf-lenti pipeline). Subsequently, a blacklist of potentially confounding nuclear mitochondrial DNA segments (NUMTs) sites was masked from the reference, as described in [2]. Finally, we used STAR to build a custom genome index for all single cell libraries (MT, GBC, GEX). Of note, MT, GEX and GBC reads are all aligned with STAR Solo – internally called both by nf-MiTo and nf-lenti – with the following command line arguments:

- `--soloType`: CB_UMI_Simple
- `-soloCBmatchwhitelist`: 1MM_multi_Nbase_pseudocounts;
- `-soloCellFilter`: Empty_Drops_CR
- `--soloUMIdedup`: 1MM_CR

which enable `EmptyDrops` CB correction and UMI deduplication, leveraging the publicly available 10x v3' CBs whitelist as reference for CBs correction.

<u>`TUNE`</u>

Given some pre-processed AFM (`nf-MiTo` step i), many hyper-parameters determine `nf-MiTo` (step ii-v) final output. To evaluate alternative hyper-parameter combinations in efficient and automated fashion, `nf-MiTo` provides the `TUNE` entry-point. This workflow allows grid-search-based hyper-parameter tuning by tracking - with a comprehensive set (n=19) of quantitative metrics, see Supplementary Information - `MiTo` toolkit – `nf-MiTo` metrics - key properties of the filtered feature space, the associated cell-cell distances, and the inferred phylogeny. These diagnostic checks enable fast and principled selection of `nf-MiTo` hyper-parameters - see Supplementary Information - `MiTo` toolkit – `nf-MiTo` hyper-parameters - prior than other computationally expensive operations – e.g., tree bootstrapping.

<u>`EXPLORE`</u>

While quantitative assessments are usually more reliable than visual inspection, direct visualization of MT-phylogenies and clones can greatly help MT-scLT analyses. Therefore, `nf-MiTo` couples the `TUNE` entry-point with the `EXPLORE` entry-point. Given a pre-processed AFM (`nf-MiTo` step i) and some associated `TUNE` output - see Supplementary Information - `MiTo` toolkit – `nf-MiTo` entry-points - the `EXPLORE` workflow produces a comprehensive set of visualizations that can be leveraged to visually explore MT-data, complementing `TUNE` quantitative metrics.

<u>`INFER`</u>

Given a pre-processed AFM (`nf-MiTo` step i) and a selected combination of hyper-parameters, the `INFER` entry-point can be used to run the complete `nf-MiTo` lineage inference pipeline (steps ii-v). This workflow includes the main data operations performed with the `TUNE` entry-point, but returns a filtered AFM, a fully annotated MT-phylogeny, and comprehensive tree metrics - see Supplementary Information - `MiTo` toolkit – `nf-MiTo` metrics. The fully annotated MT-phylogeny includes bootstrap supports [3], and - optionally, only for MT-scLT systems - MT-clones, inferred by the dedicated MiTo algorithm – see Materials and Methods – Computational Methods – MiTo – Clonal inference.

BENCH

Given a pre-processed AFM (nf-MiTo step i-iv), the BENCH entry-point run the lineage inference step of nf-MiTo (step v) with one of the 4 clonal inference methods benchmarked in this work: i) MiTo; ii) vireoSNP; iii) leiden; and iv) CClone. See Supplementary Information - MiTo benchmark - Clonal inference.

**nf-MiTo hyper-parameters**.

Each nf-MiTo entry-point - described in Supplementary Materials and Methods: nf-MiTo entrypoints - implements only a subset of nf-MiTo functionalities. On the contrary, the main nf-MiTo entry-point runs the entire pipeline (step i-v). Whichever the case, nf-MiTo capabilities can be effectively tuned by n=53 hyper-parameters. The following list includes all nf-MiTo hyper-parameters (grouped according to their pipeline step):

Step i (raw MT-reads pre-processing)

1. --pp_method: the raw MT-reads pre-processing method. Default: *maegatk*. Options: *maegatk, MiTo, cellsnp-lite, samtools, freebayes*.
2. --raw_data_input_type: see Supplementary Materials and Methods: nf-MiTo entrypoints: PREPROCESS. Default: *"fastq"*. Options: *"fastq, MAESTER"* and *"mitobam"* . This parameter is active only when --scLT_system is: MAESTER.
3. --raw_data_input: path to either the raw sequencing libraries (GEX, MT) sample sheet, or the aligned sample sheet (both in .csv format). Default: null. This parameter is active only when --scLT_system is MAESTER.
4. --afm_input: AFM assemby and processing can be done externally to nf-MiTo. The resulting (properly formatted) AFMs can be directly fed to nf-MiTo BENCH, TUNE, INFER, and EXPLORE entry-points. This argument points to a .csv file with all AFM paths (i.e., it is an AFM sample sheet) to analyze, and it is used by all entrypoints that do not involve raw MT-reads pre-processing. Default: null.
5. --ref: path to STAR index used for MT and GEX read mapping. Default: null.
6. --string_MT: MT-chromosome string identifier. Default: ChrM.
7. --whitelist: path to 10x CB whitelist used for STAR Solo CB error correction. Default: null

8. `--CBs_chunk_size`: size of the CB partition into which aligned MT-consensus reads need to be split for parallel processing (e.g., consensus sequence generation). Default: 2500. Options: [0,+inf]
9. `--fgbio_UMI_consensus_mode`: `fgbio GroupReadsByUmi --strategy` command line argument. Controls how single-cell reads are grouped by UMI sequence similarity - and mapping position - to generate read groups that will be collapsed into unique consensus reads. Default: *identity*. Options: *identity, edits, adjacency*
10. `--fgbio_UMI_consensus_edits: fgbio GroupReadsByUmi --edits` command line argument. Controls how reads are grouped by their UMI sequence similarity (i.e., the maximum number differences in UMI sequence for reads to be considered PCR duplicates of the same original RNA molecule. Default: 0. Options: 1,2,3,… .
11. `--fgbio_min_reads_mito: fgbio CallMolecularConsensusReads --min-reads` command line argument. Controls which UMI-tagged read group has enough PCR replicates (i.e., n of reads) for consensus read generation. Default: 3. Options: 1,2,3,… .
12. `--fgbio_base_quality: fgbio CallMolecularConsensusReads --min-input-base-quality` command line argument. Controls individual consensus base generation - each consensus base is generated only from raw input bases with base quality >= `--min-input-base-quality` – and `MiTo` pre-processing pile-up – only consensus bases with base quality >= `--min-input-base-quality` are counted in the final allelic tables. Default: 30. Options: any integer value.
13. `--fgbio_base_error_rate_mito:` controls maximum error rate – 1 – the fraction of raw bases that are concordant with the consensus base at some read position – allowed for a consensus base mapping to the MT-genome to be included in `MiTo` pre-processing allelic tables. Default: 0.25; Options: [0,1].
14. `--fgbio_min_alignment_quality:` controls `MiTo` pre-processing pile-up - only consensus bases from reads with alignment quality >= --`fgbio_min_alignment_quality` are counted in the final allelic tables.
15. `--path_meta:` path to cell metadata file (.csv format). This file stores additional, context specific cell metadata to include in the processed Allele Frequency Matrix. Default: null.
16. `--min_n_UMIs:` the minimum number of UMIs (GEX library) for a good quality (GEX) cell. Default: 500. Options: [0,+inf]. This parameter is active only when `–raw_input_data_type` is: fastq.
17. `--min_n_genes:` the minimum number of genes (GEX library) for a good quality (GEX) cell. Default: 250. Options: [0,+inf]. This parameter is active only when `–raw_input_data_type` is: fastq.

18. --max_perc_mt: the maximum fraction of MT counts in the GEX library allowed for a good quality (GEX) cell. Default: 0.15. Options: [0,1]. This parameter is active only when -raw_input_data_type is: fastq.
19. --n_mads: the number of Median Absolute Deviations used for GEX cell QC. Default: 3. Options: 1,2,3,5,… .

Step ii (AFM filtering, i.e., cell and MT-SNV filtering operations)

20. --cell_filter: the cell filter used to exclude low quality (MT) cells. Default: filter2. Options: filter1, filter2, nofilter.
21. --filter_dbs: whether putative MT-SNV are filtered with dbSNP and REDIdb databases. Default: true. Options: true, false.
22. --filtering: main MT-SNV filter. Default: MiTo. Options: baseline, MiTo, MQuad, miller2022, weng2024.
23. --filter_moran: enable filtering of MT-SNVs that show significant spatial auto-correlation with cell-cell distances in the MT-SNV space. Default: true. Options: true, false.
24. --min_cell_number: works with --lineage_column. Filter only categories in --lineage_column with more than --min_cell_number cells. Default: 0. Options: [0,+inf].
25. --min_cov: minimum average MT-genome coverage for a candidate MT-SNV site. Default: 5. Options: [0,+inf].
26. --min_var_quality: minimum average base calling quality for a candidate MT-SNV. Default: 5. Options: [0,+inf].
27. --min_frac_negative: minimum fraction of negative cells for a candidate MT-SNV. Default: 0.2. Options: [0,1].
28. --min_n_positive: minimum number of positive (AF>0) cells for a candidate MT-SNV. Default: 0.2. Options: [0,1].
29. --af_confident_detection: the allelic frequency at which an event of MT-SNV detection is considered “confident”. Default: 0.02. Options: [0,1]
30. --min_n_confidently_detected: the minimum number of confident detection events for a candidate MT-SNV. Default: 5. Options: [0,+inf]
31. --min_mean_AD_in_positives: the minimum average number of UMI counts for the alternative allele (AD) in positive cells (AF>0), for a candidate MT-SNV. Default: 1.25. Options: [0,+inf]

32. `--min_mean_DP_in_positives:` the minimum average number of total UMI counts in positive cells (AF>0), for a candidate MT-SNV. Default: 25. Options: [0,+inf]

Step iii (MT-genotyping)

33. `--t_prob:` value for `MiTo` genotyping probability thresholding. Default: 0.7. Options: [0,1].
34. `--min_AD:` minimum number of UMI counts for the alternative allele (AD) to assign the mutated genotype. Default: 2. Options: [1,+inf]
35. `--min_cell_prevalence:` minimum prevalence (fraction of positive cells) for a MT-SNV to be genotyped with the probabilistic approach developed in `MiTo`. Default: 0.05. Options: [0,1]
36. `--t_vanilla:` minimum allelic frequency to assign the mutated genotype using the `vanilla` genotyping method. Default: 0. Options: [0,+inf]
37. `--bin_method:` genotyping method. Default: `MiTo`. Options: `MiTo, MiTo_smooth, vanilla`.
38. `--k:` k neighbors for `MiTo_smooth` probabilistic smoothing. Default: 5. Options: [1,+inf]
39. `--gamma:` `MiTo_smooth` probabilistic smoothing weigth. Default: 0.2. Options: [0,1]
40. `--min_n_var:` minimum number of MT-SNVs to retain a cell after AFM filtering and genotyping. Default: 1. Options: [1,+inf]

Step iv (distances in MT-SNV space)

41. `--distance_metric:` the distance metric used to calculate cell-cell distances in MT-SNV space. Default: weighted_jaccard. Options: weighted_jaccard, weighted_hamming, scipy and sklearn compatible metrics and callables.

Step v (lineage inference)

42. `--tree_algorithm:` algorithm for phylogeny inference. Default: `cassiopeia.` Options: `cassiopeia, mpboot, iqtree.`
43. `--cassiopeia_solver:` tree solver from cassiopeia. Default: UPMGA. Options: UPMGA, NJ, spectral, max_cut, greedy, shared_muts.
44. `--n_boot_replicates:` number of bootstrap replicates (character matrices, distance matrices and trees). Default: 100. Options: [1,+inf]
45. `--boot_strategy:` strategy for character bootstrapping. Default: feature_resampling. Options: feature_resampling, jacknife, counts_resampling.
46. `--frac_char_resampling:` fraction of resampled characters for each bootstrapping sample. Default: 0.8. Options: [0,1]

47. `--support_method:` bootstrap support method. Default: tbe (i.e., Transfer Bootstrap Expectations […]). Options: tbe, fbp (Falsestein's Bootstrap Proportions).
48. `--annotate_tree:` annotate the inferred phylogenies with leaves and internal node statistics. Performs (MT-SNV based) clonal inference. Available only if `-scLT_method` is Redeem or MAESTER. Default: true. Options: true, false.
49. `--max_fractions_unassigned:`

Miscellanea:

50. `--scLT_system:` the scLT system from which allelic tables come from. Default: *MAESTER*. Options: *MAESTER, scWGS, RedeeM, Cas9*.
51. `--output_folder:` path to the output folder. Default: null.
52. `--lineage_column:` categorical column in AFM metadata. If different from null, it can be used to compute different statistics – e.g., MT-SNVs enrichment in each `lineage_column` category, and concordance with `MiTo` clone labels. Default: null.
53. `--K:` if `lineage_column` is not null, `--K` specifies for the number of k neighbors used to calculate kBET […] and kNN purity statistics, using `--lineage_column` labels as categorical labels. Default: 10. Options: [1,+inf]

**`nf-MiTo` metrics**

`nf-MiTo` not only provides remarkable flexibility in terms of hyper-parameter configurations, but also includes n=19 quantitative metrics to assist the user in the exploration of MT-data. The following list includes all `nf-MiTo` metrics (grouped considering their diagnostic assessment):

Mut Quality:

1. `n dbSNP`: number of filtered (i.e., after baseline and `MiTo` filter) MT-SNVs flagged as common mutation event in the dbSNP database: https://ngdc.cncb.ac.cn/databasecommons/database/id/1622
2. `n REDIdb`: number of filtered MT-SNVs flagged as common RNA-editing event in the REDIdb database: http://srv00.recas.ba.infn.it/redidb/
3. `transition vs transversion ratio`: used to quantify the deviation from the expected MT-mutational signature (i.e., transitions >> transversion)

Clonal inference accuracy:

4. `% clonal biased MT-SNVs`: % of filtered MT-SNVs that is significantly enriched (Fisher's exact test, FDR <= 0.05) within at least one lentiviral clone
5. `AUPRC`: Area Under Precision Recall Curve as described in [4]

6. `ARI`: Adjusted Rand Index, to quantify concordance between GBC labels and inferred MT-clones (see MiTo tree annotator)
7. `NMI`: Normalized Mutual Information score, to quantify concordance between GBC labels and inferred MT-clones (see MiTo tree annotator)

Tree structure:

8. `CI`: mean Consistency Index of tree characters. Ths measure assessing how well a cofinguration of character is explained by a phylogenetic tree assuming minimal evolutionary changes (i.e., Camin-Sokal parsimony)
9. `corr`: Tree- vs char- based distance correlation: Pearson's correlation between tree-based (i.e., minimum number of nodes connecting two leaves) and character-based (i.e., throughout this work jaccard distance between cell-cell MT-SNVs genotypes) cell-cell distances.

Cell connectedness in MT-SNV space:

10. `Density`: % of non-zero entries in the binarized AFM
11. `Transitivity`: transitivity (i.e., clustering coefficient) of the cells shared-MT-SNVs graph
12. `Mean path length`: average cell-cell path-length on the cells shared-MT-SNVs graph
13. `Average degree`: average cell degree across the cells shared-MT-SNVs graph
14. `LCC`: largest connected component of the cells shared-MT-SNVs graph

Genotype variation:

15. `Haplotype redundancy`: % of unique MT-haplotypes (i.e., beared by single-cells) considering all MT-hatplotypes observed in a population of cells
16. `Median n of MT-SNVs per cell`

Cellular Yield:

17. `n_GBC_groups:` number of GBC clones
18. `n_cells:` number of filtered cells
19. `n_vars:` number of filtered MT-SNVs

**Data generation**

This section provides technical details about all the experimental work needed to generate our benchmarking datasets. For a general overview, see Materials and Methods – Data generation.

**Dual scLT experiments**

Perturb-seq (GBC) library preparation and sequencing

A semi-nested PCR approach was used to ensure maximum yield. This PCR employs two specific primers (targeting the adapter sequences inserted during the GEX library production, one couple for each sample), and one reverse non-specific primer (annealing to a constant portion of the BFP sequence), which is constant in all barcode fragments. Presence of the specific ~400 bp fragment was assessed for each sample using 2% agarose (Canvax Biotech) gel electrophoresis. The PCR product was diluted 1:1000 to reduce the amount of other non-specific fragments. Then, we performed a 2nd reaction with the setting and primers reported in Suppl. Table 1. The presence of the diagnostic ~400bp product was checked for every sample using 2% agarose (Canvax Biotech) gel electrophoresis. PCR products were purified with QIAquick PCR purification kit (Qiagen, see […] et al., 2023 for the primer sequences and the PCR cycles). GBC libraries (2ng per sample) were sequenced with the NovaSeq 6000 Sequencing System (~10k reads per cell, ~60M reads per sample).

MAESTER (MT) library preparation and sequencing

The mitochondrial transcripts (MT-) library was generated from 10x barcoded cDNA following the original MAESTER protocol. MAESTER, MT-transcripts-targeting primers are used together with P5 and P7 adapter-annealing primers. Two PCRs are needed (i.e., PCR1 and PCR2). For PCR1, 240 ng of 10x cDNA were amplified in 12 parallel reactions (20 ng of cDNA per reaction), each targeting a specific region of the MT-genome (through region-specific primer mixes). These PCRs (12 per sample +1 negative control) were performed in 96-well plates, with the following set up:

| **Reagents** | **Volume (ul)** | **concentration** |
|---|---|---|
| cDNA + H20 | 15 | 20ng |
| Primer P5 | 1 | 10uM |
| Mitochondrial primer Mix | 4 | 1uM |
| KAPA Hifi | 20 | |
| Tot volume | 40 | |

| **Reaction** | **Temperature** | **Time** | **Cycles** |
|---|---|---|---|
| Initial denaturation | 95 °C | 3min | X 1 |
| | 98 °C | 20 sec | X 6 |
| | 65 °C | 15 sec | |

| | | | |
|---|---|---|---|
| | 72 °C | 3min | |
| Final extension | 72 °C | 5min | X 1 |

Amplicons of interest were captured with the AMPure Bead kit, with a 0.8x ratio, followed by PCR2:

| **Reagents** | **Volume (ul)** | **concentration** |
|---|---|---|
| cDNA amplified (PCR1) | 18 | |
| Primer P5 | 1 | 5uM |
| Primer P7 | 4 | 5uM |
| KAPA Hifi | 20 | |
| Tot volume | 40 | |

| **Reaction** | **Temperature** | **Time** | **Cycles** |
|---|---|---|---|
| Initial denaturation | 95 °C | 3min | X 1 |
| | 98 °C | 20 sec | X 6 |
| | 60 °C | 30 sec | |
| | 72 °C | 3min | |
| Final extension | 72 °C | 5min | X 1 |

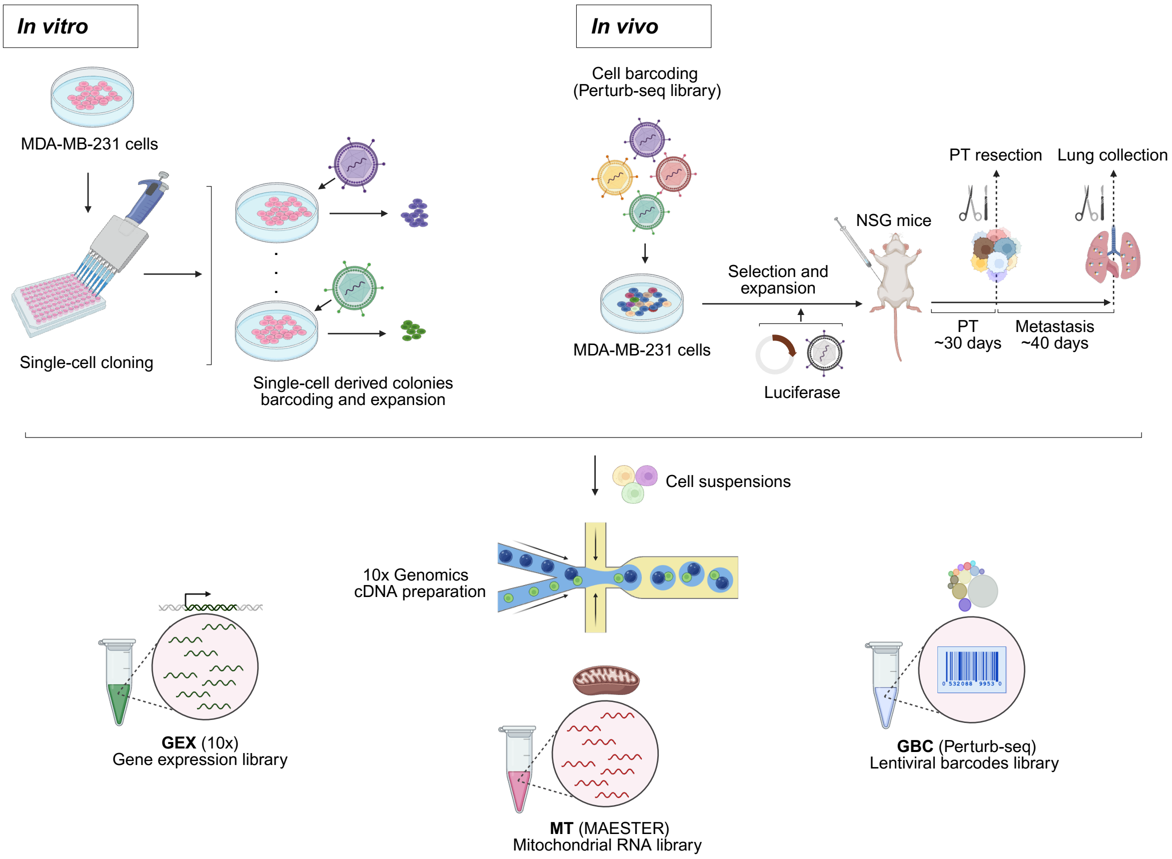


**Supp. Fig. 1. Dual-lineage tracing experimental design and generation of benchmarking datasets. Left panel:** Schematic of the *in vitro* clonal mixture model (MDA_clones). Single MDA-MB-231 cells were isolated by FACS and expanded to generate clonal colonies. After ~30 days, each colony was transduced with a unique barcode-expressing lentivirus. Eight barcoded colonies were then mixed in defined proportions to form a controlled clonal mixture. **Right panel:** Schematic of the *in vivo* longitudinal model. MDA-MB-231 cells transduced with a high-complexity Perturb-seq lentiviral barcode library were orthotopically injected into immunocompromised NSG mice. After ~30 days of primary tumor (PT) growth, tumors were resected, followed by ~30 additional days to allow formation of lung metastases (Mets). One matched PT–lung pair was selected for further profiling (MDA_PT and MDA_lung). **Bottom panel:** All specimens underwent 10x Genomics scRNA-seq. A modified MAESTER protocol enabled simultaneous capture of gene expression (GEX), mitochondrial variants (MTs), and lentiviral barcodes (GBCs), generating three high-quality, multi-modal benchmarking datasets.

#### `MiTo` benchmark

This section provides details about `nf-MiTo` hyper-parameter tuning - and therefore, choice of default values for `MiTo` functions' arguments. Before dealing with this section, see Supplementary Information – `MiTo` toolkit - `nf-MiTo` hyper-parameters, and Supplementary Information – `MiTo` toolkit - `nf-MiTo` metrics for reference.

**Preliminary assessment**

After multiple rounds of development and testing for each step of the `nf-MiTo` pipeline – we set the 0-shot hyper-parameter configuration of `nf-MiTo` based on clonal inference accuracy – `ARI`, `NMI`, `AUPRC`, `% clonal biased MT-SNVs` - and cellular yield – `n_GBC_groups`, `n_cells`, `n_vars` - evaluations. Hereafter, we will use the term "clonal inference performance" to indicate joint optimization of clonal inference accuracy and cellular yield metrics. During this phase, we recognized 5 hyper-parameters that highly influenced clonal inference performance:

i. `--cell_filter`: the MT-based cell filter, based on MT-genome coverage statistics. See Materials and Methods – Computational Methods – `MiTo` – Cell filtering.
ii. `--af_confident_detection`: the allelic frequency of confident MT-SNV detection.
iii. `--min_n_confidently_detected`: the minimum number of confidently detected events to filter a MT-SNV
iv. `--min_mean_AD_in_positives`: the minimum number of alternative allele (AD) UMI counts in +cells (AF>0) to filter a MT-SNV. See Materials and Methods – Computational Methods – `MiTo` – MT-SNVs filtering.
v. `--min_AD`: the minimum AD counts needed to assign alternative genotypes. See Materials and Methods – Computational Methods – `MiTo` – MT-genotyping.

To fine-tune these 5 hyper-parameters, we tested - with `nf-MiTo TUNE` - 90 unique combinations of these hyper-parameter values across our 3 benchmarking datasets – leaving other one unchanged. Supp. Fig. 2a highlights two major trends. First, clonal inference on the MDA_clones and MDA_lung datasets is consistently easier than on the MDA_PT dataset. Second, especially considering the MDA_PT dataset clonal inference accuracy and cellular yield are often anti-correlated: stringent hyper-parameter combinations yield simple MT-SNV spaces - few cells and GBC clones - making clonal inference easier, but loosing a substantial portion of the data, while permissive ones yield the opposite results. This trend implies that optimal AFM filtering should clean up the data just enough to let the real phylogenetic signal emerge without over-cleaning the data. All tuned hyper-parameters have an impact on this trade-off, but `--cell_filter` represent a clear example, as shown in Supp. Fig. 2b.

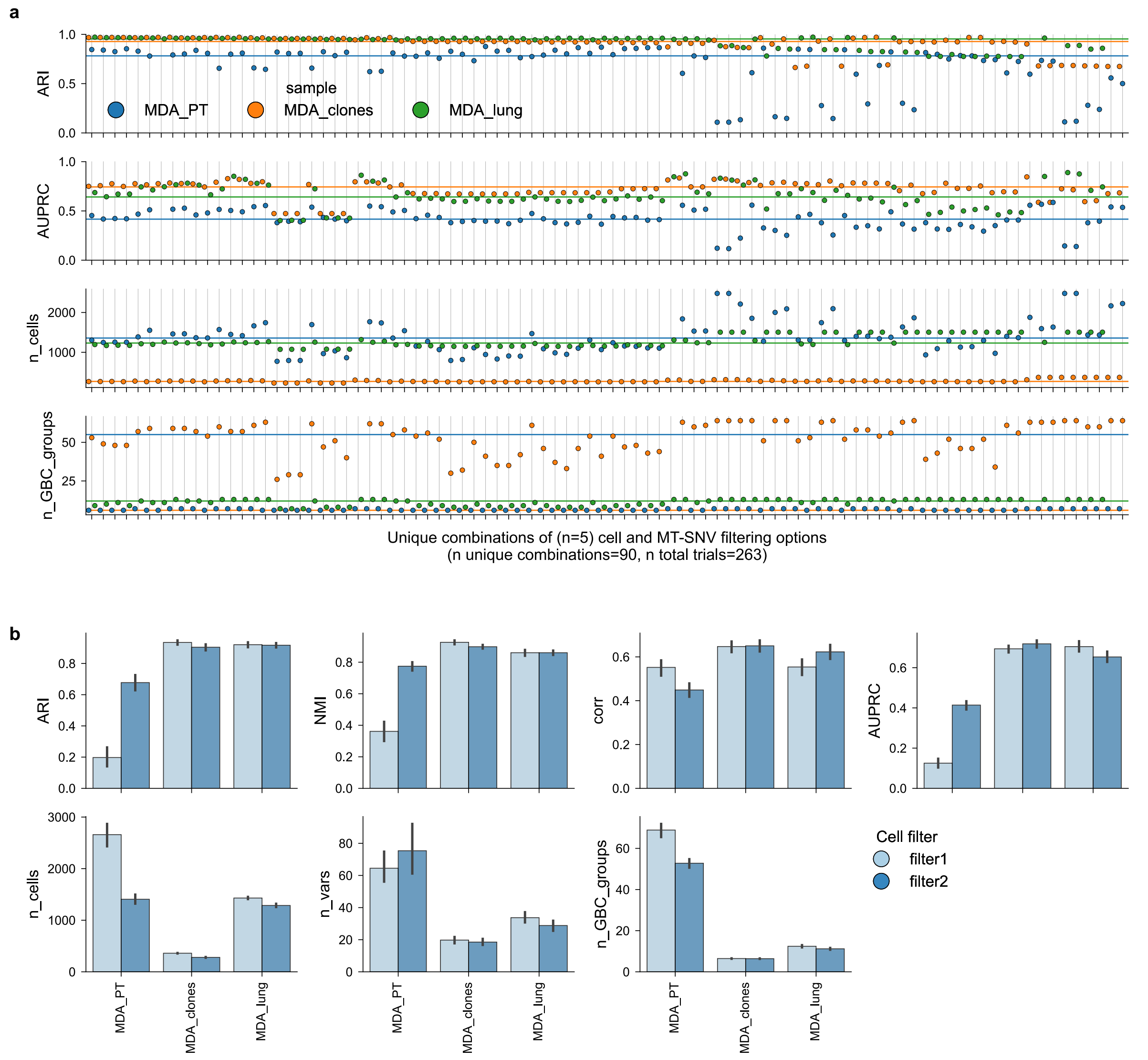


**Supp. Fig. 2. Preliminary assessments of lineage inference performance. a.** General overview of clonal inference performance – ARI, AUPRC – and cellular yield metrics – n_cells, n_GBC_groups – for n=90 unique combinations of 5 `nf-MiTo` hyper-parameters – see Supplementary Information – `MiTo` toolkit - `nf-MiTo` hyper-parameters and Supplementary Information – `MiTo` toolkit – `nf-MiTo` metrics for reference. x-axis: unique combinations; y-axis: `nf-MiTo` metric. Dots represent metric values for a specific combination and sample (see legend). **b.** `nf-MiTo` metric values across unique hyper-parameters combinations - same as in **a** – grouped by sample and `MiTo` cell filter.

#### Cell filtering

To systematically evaluate the impact of cell filtering strategies on lineage inference performance, we perturbed the `--cell_filter` and `--min_cell_number` hyper-parameters for the lowest and highest complexity dataset - i.e., MDA_clones and MDA_PT, respectively. See also Materials and Methods - Computational Methods – `MiTo` – Cell filtering.

Considering MDA_clones, Supp. Fig. 3a-c shows increased loss of cells and quality of MT-genome coverage statistics with progressively stringent cell filtering strategies – i.e., No filter, filter1 and filter2. Combined with different `--min_cell_number` values – i.e., including GBC clones with more than 0 or 5 cells - Supp. Fig. 3d did not dramatically alter the clonal composition of the dataset, consistent with the small number of balanced clones of this sample. However, these effects are much more dramatic considering MDA_PT as highlighted by Supp. Fig. 3e-h, giving the high number of low-prevalence clones in this sample.

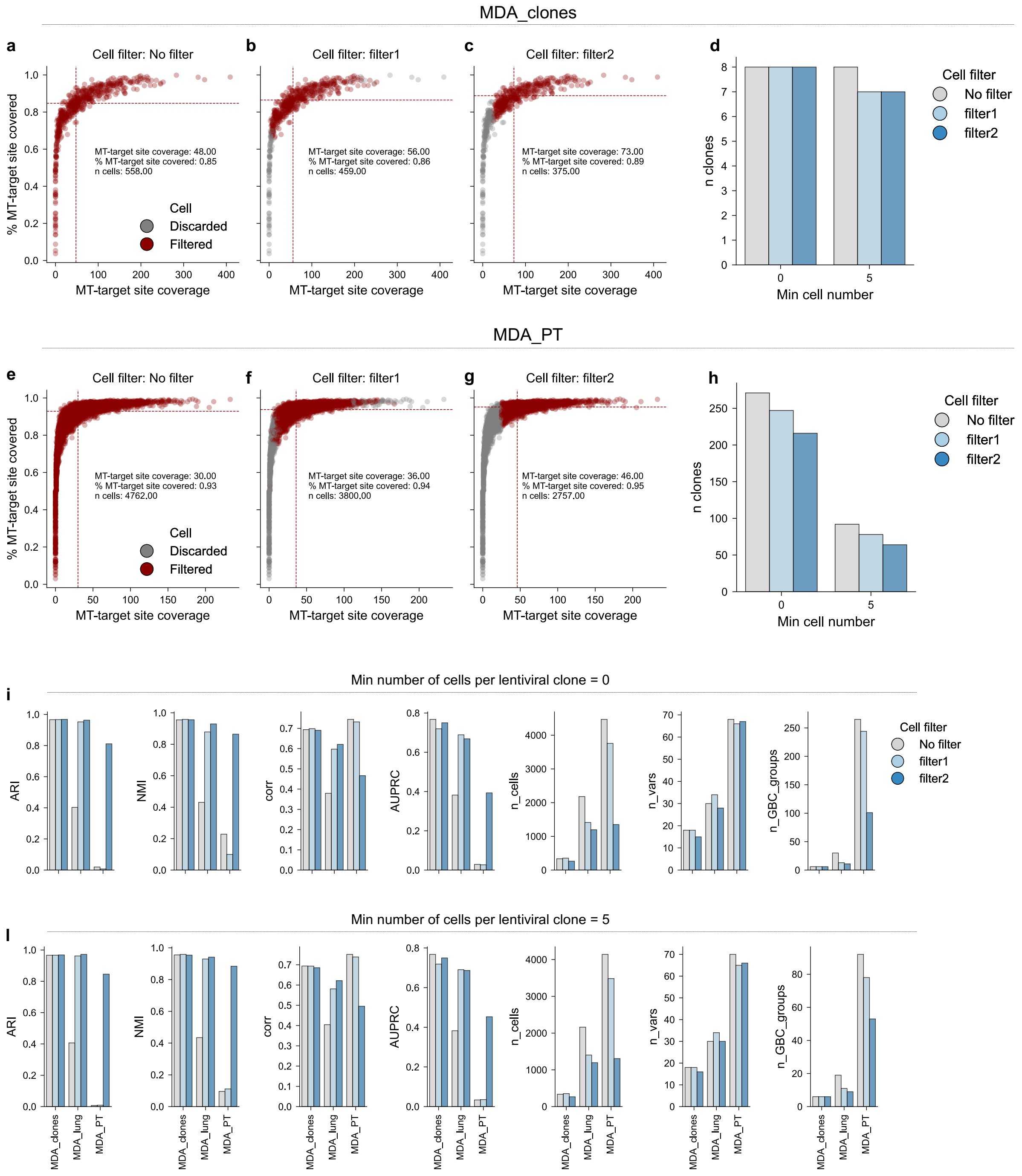

**Supp. Fig. 3. Cell filtering benchmark. a-c.** Same scatterplot shown in **Fig. 2d**, showing cell-specific statistics of MT-coverage used for cell filtering in the MDA_clones dataset. The 3 panels depict cells filtered (and filtered out) in case with no filtering (**a**), "filter1" (**b**) and "filter2" (**c**) – see Materials and Methods - Computational Methods – `MiTo` – Cell filtering. **d**. Number of clones in MDA_clones dataset after cell filtering and additional removal of cells from GBC clones with <0 or 5 cells - i.e., min_cell_number, see Supplementary Information – `MiTo` toolkit – `nf-MiTo` hyper-parameters. **e-h.** Same as in **a-d,** but for the MDA_PT dataset**. i.** Selected Clonal inference accuracy, Cellular yield and Tree structure metrics for the across samples, grouped by cell filter, without downstream removal of any clone low prevalence GBC-clone – i.e., min_cell_number = 0. **j.** Same as in **i**, but with min_cell_number = 5.

Clonal inference accuracy, Cellular yield and Tree structure metrics – see Supplementary Information – `MiTo` toolkit - `nf-MiTo` metrics – exhibit substantial variation across all 3 datasets – MDA_clones, MDA_PT, MDA_lung - and filtering strategies **Supp. Fig. 3i-l**, with MDA_lung representing a middle situation between MDA_clones and MDA_PT. Progressively stringent cell filtering strategies dramatically reduced the number of cells and GBC clones for both MDA_lung and MDA_PT samples, but this reduction was not mirrored by a similar MT-SNV losses. Consistently, clonal inference accuracy metrics dramatically improved after low-quality cell removal, with the most evident effect on the MDA_PT dataset. The correlation between tree-based and character-based cell-cell distances varied unexpectedly, increasing with cell filtering stringency for MDA_lung but decreasing for MDA_PT, and remaining substantially identical for MDA_clones. Reassuringly, regardless of MT-coverage-based cell filtering strategies, including very low-prevalence clones – 0-5 cells – did not alter clonal inference accuracy.

In summary, these data demonstrated the fundamental importance of cell filtering based of MT-coverage metrics. This analytical step is mandatory especially in high clonal complexity datasets, where only the highest stringency `MiTo` cell filter – filter2 – provided AFMs with identifiable GBC clones.

**Raw reads pre-processing and AFM filtering**

Once low-quality cells have been discarded, the `nf-MiTo` filters high-confidence MT-SNVs – see Materials and Methods - Computational Methods – `MiTo` – MT-SNVs filtering. Several approaches has been proposed for this task, with raw reads pre-processing – collection of alternative and reference (or total) read/UMI counts for some MT-genome position – and MT-SNV filtering – filtering of high-confidence variant sites - performed jointly, or in separate stages. Here, we have benchmarked lineage inference performance of 5 pre-processing pipelines combined with 2 MT-SNVs filters.

Among pre-processing pipelines – the `--pp_method` hyper-parameter, supporting `maegatk`, `MiTo`, `cellsnp-lite`, `samtools` and `freebayes` options – we first included the 2 state-of-the-art pipelines for expressed MT-SNVs pre-processing – maegatk and cellsnp-lite – and subsequently added 2 other pipelines developed for bulk-level nuclear SNVs calling – samtools and freebayes - re-adapted to single-cell MT-SNVs calling, as done in […]. As illustrated in Supp. Fig. 4, all of these tools work with different working principles.

Specifically, maegatk leverages 10x Unique Molecular Identifiers (UMIs) to group PCR replicates into distinct molecular groups and collapse them into a single "consensus" sequence. These sequences are then re-mapped to the MT-genome, and custom criteria are used to build cell-specific pile-up tables, where extensive summary statistics are registered – e.g., consensus UMI counts and mean base calling quality for each allele (A, C, T, G), strand orientation (forward and reverse), and MT-genome position. maegatk AFMs are assembled from these pile-up tables. Importantly, since maegatk returns both (cell-specific) MT-genome site coverage, all cell filtering strategies benchmarked in Supplementary Information – MiTo benchmark – Cell filtering can be used to filter good-quality cells. Since maegatk returns statistics for all MT-genome site, MT-SNVs filtering needs to be performed in a separate step. As, previously shown [5], maegatk pile-up tables can processed to compute MT-SNV-specific summary statistics to inform MT-SNVs selection – which is usually performed with post-hoc thresholding.

On the other hand, cellsnp-lite uses 10x UMIs and bulk-level summary statistics to call an initial, large subset of MT-SNVs, returning alternative and reference alleles UMI counts. Different form maegatk, cellsnp-lite output does not include MT-genome site coverage - preventing downstream cell filtering, at least based on this property. Moreover, cellsnp-lite output does not include allele-specific mean base-calling quality – preventing maegatk-like downstream MT-SNVs filtering. Since cellsnp-lite variant calling still employs very loose MT-SNV filters, the retuned MT-SNVs call-set can be filtered further with the companion MQuad approach [6] (see below).

Differently from the maegatk and cellsnp-lite, samtools and freebayes directly use 10x sequencing reads (i.e., no UMIs) for allele counting. These two latter tools can take single-cell MT-libraries and jointly perform variant filtering – with proprietary statistical frameworks – and reference/alternative alleles pile-up. As for cellsnp-lite, samtools and freebayes output does not allow direct - MT-coverage-based - cell filtering. However, their built-in MT-SNVs filters already provide conservative MT-SNV call-sets, preventing the need for other downstream MT-SNVs filtering.

| Pipeline | UMI-based | Consensus calling | Returns coverage | Returns quality | Downstream cell filters | Buil-in mut filter | Downstream mut filters |
|---|---|---|---|---|---|---|---|
| maegatk | yes | yes | yes | yes | filter1, filter2 | No | MiTo, MQuad |
| MiTo | yes | yes | yes | yes | filter1, filter2 | No | MiTo, MQuad |
| cellsnp-lite | yes | no | no | no | No filter (or external cell subset) | Yes | MQuad |
| samtools | no | no | no | no | No filter (or external cell subset) | Yes | baseline (after built-in) |
| freebayes | no | no | no | no | No filter (or external cell subset) | Yes | baseline (after built-in) |

**Supp. Fig. 4. Raw MT-reads pre-processing pipelines.** Summary tables including the most relevant properties of the 5 pre-processing pipelines benchmarked.

To test whether additional tuning of `maegatk` error correction approach could boost lineage inference performance, we developed the `MiTo` pre-processing pipeline. The core methodology of this latter pipeline is extremely similar to `maegatk`, apart from some key modifications. In summary, `MiTo` pre-processing performs more stringent base calling than previously developed in the original `maegatk`. Specifically, the original `maegatk` implementation uses `fgbio CallMolecularConsensus`[7] for consensus sequence generation, *without* filtering any base at this step - all observed bases in a group of reads assigned to the same UMI are used to generate the consensus sequence. In addition, `fgbio CallMolecularConsensus` produces unaligned reads with tags annotating two base-specific consensus metrics: i) cd*:* the "consensus depth" or "UMI group size", the number of good quality reads that were used for fgbio sequence consensus, and ii) ce*:* the "consensus error", the number of discordant bases from the one registered as final consensus base. In its original implementation, `maegatk` enable filtering consensus reads and bases – to register them into its pile-up tables -, but does not consider base-specific UMI-group size and consensus error as additional indicators of reliable molecular information. Since recent work attempted to make single-cell MT-SNV calls with single-molecule detection[8] - i.e., basecalls, supported by a single consensus UMI - we hypothesized that even a small fraction of low-quality/weakly-supported consensus UMIs could lead to sub-optimal lineage inference accuracy. To test this hypothesis, `MiTo` pre-processing: i) uses only Q30 bases for consensus sequence calling - `fgbio CallMolecularConsensus --min-input-base-quality` parameter - and ii) include filtering of individual consensus bases according to their UMI group size and consensus error.

This modifications produced very similar outputs overall, as shown in Supp. Fig. 5-6. Considering the MDA_clones sample, the MT-cell coverage was extremely similar across pipelines, with `MiTo` pre-processing - compared to `maegatk` - yielding fewer counts (median 66 vs 80 UMI counts across all MT-genome sites, Supp. Fig. 5a, and 74 vs 89 UMI counts across MAESTER target sites, Supp. Fig. 6), a smaller fraction of target sites covered (median 85% vs 89%, Supp. Fig. 5b), and smaller UMI group sizes (median 10.2 vs 10.5, Supp. Fig. 5c). However, `maegatk` recorded ~83k basecalls with average consensus score <0.7 - i.e., 1-consensus error of supporting consensus UMIs, 1.81% of total consensus basecalls, Supp.

Fig. 5e - and ~120k basecalls with average base-calling quality <30 - 2.61% of total consensus basecalls, Supp. Fig. 5d - that were absent in `MiTo` preprocessing base-calls. Strikingly (Supp. Fig. 5f), these apparently minor differences resulted in dramatically different (i.e., ~3-fold difference) numbers of variant basecalls - raw basecalls with at least one UMI supporting an alternative allele for some cell at some MT-genome position - with median 142 (+-71) and 442 (+-160) variant basecalls across cell for `MiTo` preprocessing and `maegatk`, respectively.

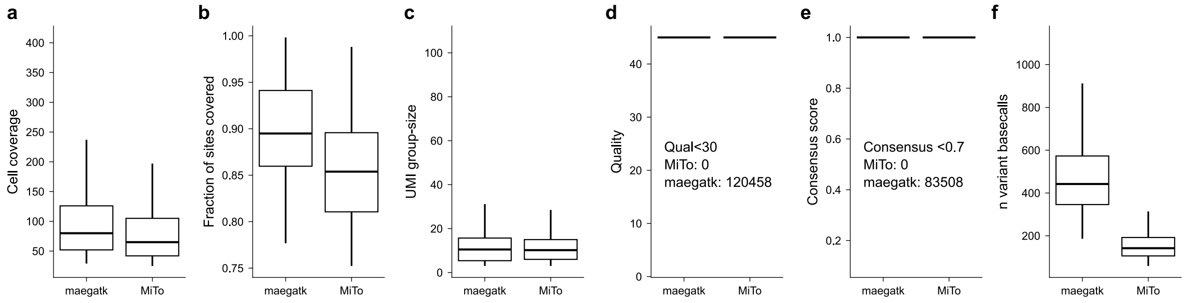


**Supp. Fig. 5. `MiTo` pre-processing vs `maegatk`.** Detailed summary statistics of `MiTo` and `maegatk` pre-processing output for the MDA_clones sample. **a.** Median (across all positions in the MT-genome) cell coverage – i.e., number of consensus UMIs per MT-genome position. **b.** Fraction of MT-genome sites covered – i.e., at least one consensus UMI – per cell. **c.** Mean consensus UMI group size – i.e., average number of reads that generated the consensus UMIs supporting a certain base in a given cell and MT-genome position. **d.** Mean base calling quality – i.e., average base calling quality of the reads that generated the consensus UMIs supporting a certain base in a given cell and MT-genome position. **e.** Mean consensus score– i.e., average consensus score of the consensus UMIs supporting a certain base in a given cell and MT-genome position. **f.** Number of variant base-calls detected per cell.

`MiTo` pre-processing and `maegatk` return very similar output formats - with `MiTo` adding mean consensus score and group size for each variant allele as additional metric. As such, as described for `maegatk`, `MiTo` requires downstream, `maegatk`-like filtering.

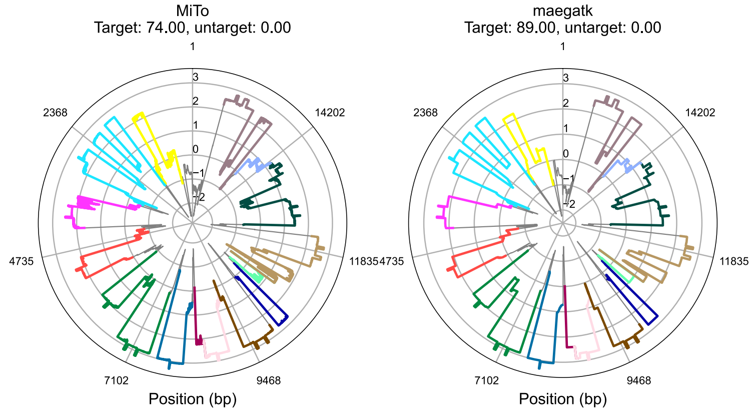

**Supp. Fig. 6. Cell coverage differences between `MiTo` and `maegatk`.** Same coverage plot shown in **Fig. 2c**, but for the MDA_clones dataset, considering `MiTo` and `maegatk` AFMs.

Different works have chosen different threshold for similar summary statistics for MT-SNVs filtering[5,8]. The only available alternative to these empirical thresholding approaches is represented by the `MQuad` method – a probabilistic approach[6] - that adaptively select the most conservative set of MT-SNVs providing enough statistical evidence for the existence of both negative and positive cell populations. `MiTo` combines different summary statistics and statistical tests for robust MT-SNVs filtering – see the `MiTo` filter in Materials and Methods – Computational Methods – `MiTo` – MT-SNVs filtering. This filter requires MT-SNV base calling quality and MT-genome site coverage, and therefore, it can only be applied to `maegatk` and `MiTo` AFMs.

In this scenario, we systematically benchmarked allowed combinations – see Supp. Fig. 4 – of pre-processing methods and MT-SNVs filters (Supp. Fig. 7). In order to eliminate potential cell-filtering biases – i.e., differences in lineage inference performance deriving from the analysis of different cell subsets within the same sample – we restricted our benchmark to a unique cell subset for every sample. To do this, for each sample AFM, we selected a single subset of good-quality cells ("filter2" cell-filtering, applied to each sample `maegatk` AFM), and manually filtered all AFMs to include only these cells. Then, we first compared the 5 pre-processing pipelines coupled to their matched MT-SNV filters – e.g., `MiTo` filter for `maegatk` and `MiTo`, `MQuad` for `cellsnp-lite` (Supp. Fig. 7). Strikingly, only `MiTo` and `maegatk` achieved good concordance between inferred- and GBC-clones - ARI and NMI > 0.8. All the other methods completely failed to recover ground-truth clones (Supp. Fig. 7a,b,d). Considering cellular yield, we observed variable numbers of cells (Supp. Fig. 7e), MT-SNVs (Supp. Fig. 7f), and GBC clones (Supp. Fig. 7g) across pipelines. This naturally followed our strict requirement according to which only cells bearing at least 1 MT-SNV can progress to lineage inference - and thus, even starting from the same initial cell subset, MT-SNVs filtering can further constrain the final number of analyzed cells.

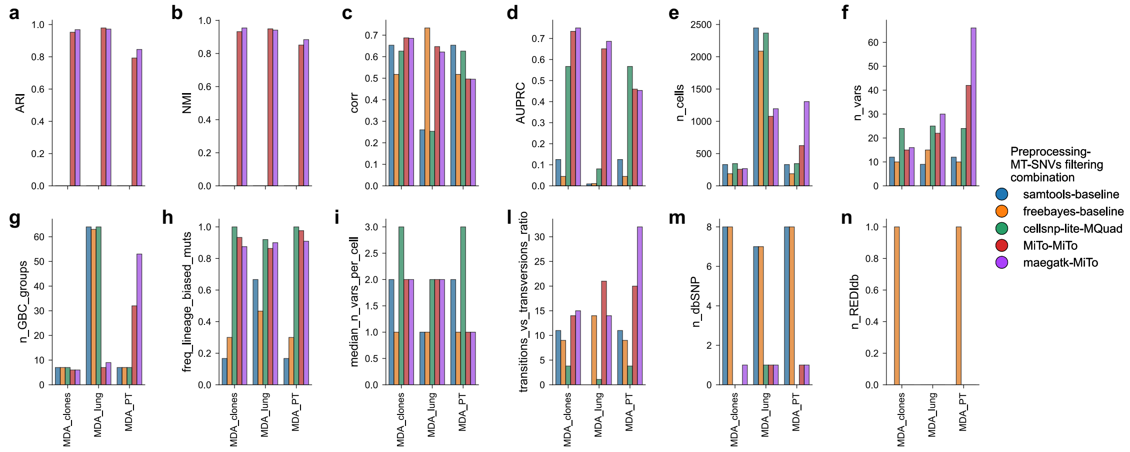


**Supp. Fig. 7. Pre-processing pipeline-MT-SNVs filter combinations benchmark.** Detailed comparison between 5 combinations of pre-processing pipelines and MT-SNVs filters. See text and Supplementary Information – `MiTo` toolkit – `nf-MiTo` metrics for individual metrics.

Interestingly, `samtools` and `freebayes` consistently filtered fewer MT-SNVs than other methods (Supp. Fig. 7f), but this did not always translate to lower number of analyzed cells – likely to high numbers of false positive events, Supp. Fig. 7e. Consistently, `samtools` and `freebayes` filtered higher numbers of un-informative MT-SNVs (Supp. Fig. 7m-n), showed lower fractions of clonally enriched mutations (Supp. Fig.7h), and lower transitions vs transversions ratios (Supp. Fig. 7l). These data highlighted poor suitability of these tools – developed for nuclear SNVs and bulk DNA sequencing data - to expressed MT-data pre-processing. On the contrary, `MiTo`, `maegatk`, and `cellsnp-lite` consistently filtered more MT-SNVs, with `maegatk` ranking first in terms of number of MT-SNVs (Supp. Fig. 7f), analyzed cells (Supp. Fig. 7e), and GBC clones (Supp. Fig. 7g), especially considering the sample with highest clonal complexity MDA_PT. Surprisingly, this higher cellular yield was not associated to lower MT-SNVs quality. In particular `maegatk` and `MiTo` – despite different stringencies in base-calling, see Supp. Fig. 5 – showed highly comparable numbers of un-informative MT-SNVs (Supp. Fig. 7m-n), fractions of clonally enriched mutations (Supp. Fig.7h), transitions vs transversions ratios (Supp. Fig. 7l) – with the latter metric even higher for `maegatk` in the MDA_PT sample. Conversely, `cellsnp-lite` performed sub-optimally compared to `maegatk` and `MiTo`, especially considering cellular yield metrics in MDA_PT (Supp. Fig. e-g), and MT-SNVs quality across all samples – poor transition vs transversion ratios (Supp. Fig. l). These data demonstrated the added value of UMI-based consensus-error correction in MT-reads pre-processing. Moreover, since increased stringency in consensus base-generation and filtering – `MiTo` vs `maegatk` – reduced cellular yield without being paralleled by increased clonal inference accuracy, this benchmark demonstrated that the original maegatk approach is sufficient to achieve optimal lineage inference performance, especially in high-complexity datasets.

Given these results, we asked whether the observed superiority of MiTo and maegatk was dependent on their downstream MiTo MT-SNV filter. To test this hypothesis, we evaluated whether their clonal inference performance was altered by swapping the MiTo filter with the MQuad filter (Supp Fig. 8).

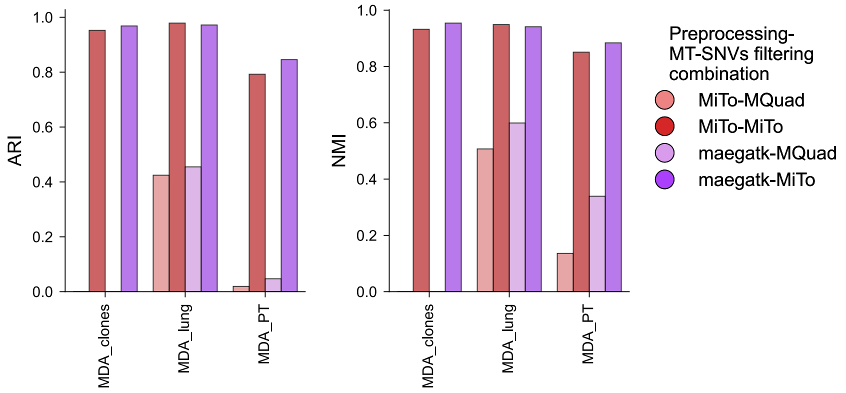


**Supp. Fig. 8. MiTo vs MQuad variant filtering strategies for maegatk and MiTo pre-processing AFMs.** Clonal inference accuracy – i.e., ARI, NMI, see Supplementary Information – MiTo toolkit – nf-MiTo metrics - comparison between MiTo and MQuad MT-SNVs filters applied on maegatk and MiTo pre-processing AFMs.

This benchmark demonstrated that MiTo and maegatk performance is tightly linked to the downstream MT-SNVs filter that is applied to their raw AFMs. Indeed, our MiTo filter is better suited to extract "informative" MT-SNVs than the state-of-the-art MQuad filter, thanks to its comprehensive, orthogonal criteria to detect confident MT-SNVs.

We than asked which was the impact of individual hyper-parameters – tuning the stringency of MiTo MT-SNVs filter. Specifically, we benchmarked n=96 combinations of 4 hyper-parameters values – see Supplementary Information – MiTo toolkit - nf-MiTo hyper-parameters:

- `--min_mean_AD_in_positives`: [ 1.00, 1.25, 1.50 ], default = 1.25;
- `--af_confident_detection`: [ 0.01, 0.02, 0.03, 0.05 ], default = 0.02;
- `--min_n_confidently_detected`: [ 2, 5 ], default = 2;
- `--filter_moran`: [ true, false ], default = true;
- `--filter_dbs`: [ true, false ], default = true;

Supp. Fig. 9 illustrates the related results. Starting from the complete set of unfiltered MT-SNVs - `maegatk` AFMs, post cell filtering Supp Fig. 9a – different filterings yielded MT-SNVs with variable contamination – i.e., variants with high-prevalence at the population level (Supp Fig. 9b, top panel) and recurrent RNA editing events (Supp Fig. 9b, bottom panel). Considering individual parameters, increasing the minimum value of (mean) AD counts in +cells (Supp Fig. 9c) to select a MT-variant resulted in higher clonal inference accuracy (ARI), but lower cellular yield (n cells, n vars). Interestingly, even small changes of this parameter value – 1.00 → 1.25 - dramatically reduced the connectedness – average degree – of cells in MT-SNV space. However, this effect was not as much dramatic for the quality of selected MT-SNV - as measured by transitions vs transversions ratio. Notably, all of these effects were more pronounced for the high-complexity dataset (MDA_PT), compared to the others. Despite having slightly less impact, the AF threshold for "confident" MT-SNV detection behave largely the same way (Supp Fig. 9d), with more stringent values giving higher clonal inference accuracy, lower number of cells, lower connectedness, with decreasing values of transitions vs transversions ratio. Compared to these two hyper-parameters, the minimum number of confident detection events had negligible effect of clonal inference accuracy and the number of cells recovered, while impacting on the number of filtered MT-SNVs (Supp Fig. 9e). On the contrary, removing downstream filters `--filter_moran` and `--filter_dbs` parameters, see Supplementary Information – `MiTo` toolkit - `nf-MiTo` hyper-parameters and Materials and Methods – Computational Methods – `MiTo` – MT-SNVs filtering – had significant impact on all reported metrics (Supp Fig. 9f), with variable effect considering different dataset. In terms of accuracy, removing MT-SNVs that did not exhibit significant spatial structure increased performance for both MDA_clones and MDA_PT, but not MDA_lung, while removing common MT-SNVs variants and RNA-edits impacted dramatically MDA_clones accuracy, but not MDA_PT and MDA_lung ones. Notably, the number of variants and cell connectedness in MT-SNV space dramatically increased disabling the spatial filter – no spatial – highlighting the importance of this criterion to remove diffused but poorly clustered variants – likely technical artifacts – that impair accurate lineage inference. Together, these data showcased the effectiveness of `MiTo` MT-SNV filter, dissecting the individual contributes of its variant selection criteria. In addition, these data argue against the feasibility of single-molecule detection (i.e., 1 UMIs) of expressed MT-SNV, but suggest that even weakly detected MT-SNVs – even only 2 detection events with AF=0.02 and mean number of RNA molecules=1.25 – can effectively contribute to recover ground-truth clonal structures, if effectively double-checked –i.e., removed un-informative variants and technical artifacts.

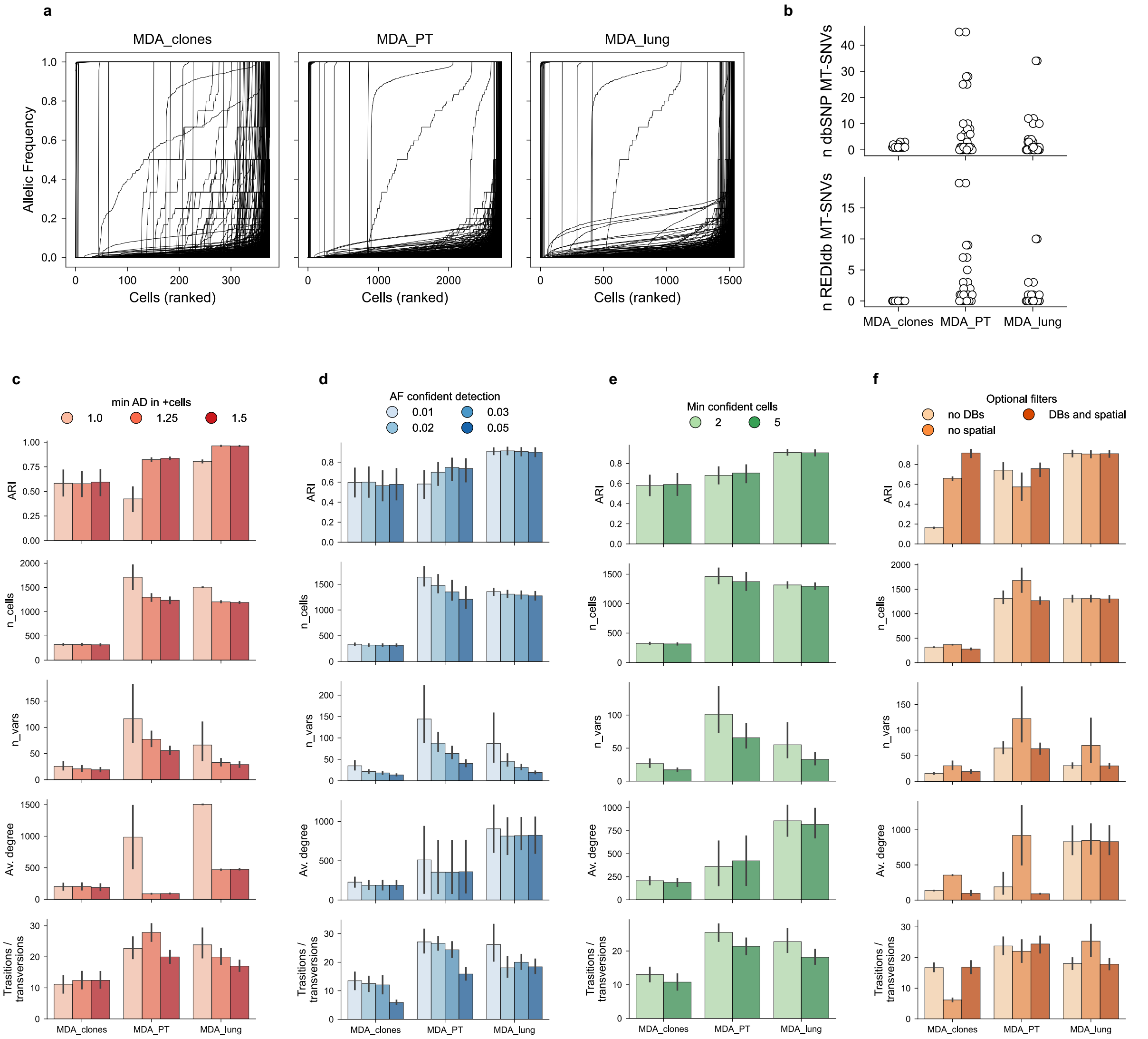


**Supp. Fig. 9. Tuning of `MiTo` MT-SNVs filter hyper-parameters. a.** Allelic frequency distribution of all detected – i.e., prior than AFM filtering - MT-SNVs in MDA_clones, MDA_PT and MDA_lung datasets. Each line represents a single variant. The x-axis represents the order of – quality-controlled – cells for each variant, while the y-axis holds AF values. **b.** Number of un-informative MT-SNVs across datasets and hyper-parameters combinations. Top: common MT-SNVs (from dbSNP). Bottom: Common MT-RNA edits (from REDIdb). **c.** Adjusted Rand Index, number of cells, number of filtered MT-SNVs, average degree of cells in their shared MT-SNVs graph, and transition vs transversion ratio across samples, varying the value for the minimum number of consensus UMIs supporting the alternative allele in positive cells (AF>0). **d.** Same metrics in **c**, varying the AF threshold to consider a MT-SNV detection event "confident". **e.** Same metrics in **c-d,** varying the minimum number of confident detection events required to filter a MT-SNV. **f.** Same metrics in **c-e,** applying different combinations of MiTo optional filters – **i.e.,** removing the "spatial" filter testing MT-SNVs significant auto-correlation with cell-cell MT-distances, and the "DBs" filter, which uses dbSNP and REDIdb databases to filter out un-informative variants. For **c-e**, see Supplementary Information – `MiTo` toolkit – `nf-MiTo` hyper-parameters and Supplementary Information – `MiTo` toolkit – `nf-MiTo` metrics.

Given robust filtering – and genotyping - of MT-SNVs, we next asked how much of them were significantly enriched – Fisher's Exact test, pvalue <= 0.05 – in ground-truth GBC clones (Supp Fig. 10). Here, we found that a substantial fraction of GBC clones is genetically defined by at least one MT-SNVs (Supp Fig. 10a). However, the median number of MT-SNVs per clone was low – 1, 2, and 3 in MDA_PT, MDA_lung and MDA_clones, see Supp Fig. 10b – and linked to the average MT-library sequencing depth (data not shown). While encouraging for clonal structure recovery – i.e., the one defined by lentiviral barcoding -, this data also pose a fundamental limit in terms of solving more recently occurred sub-clonal events.

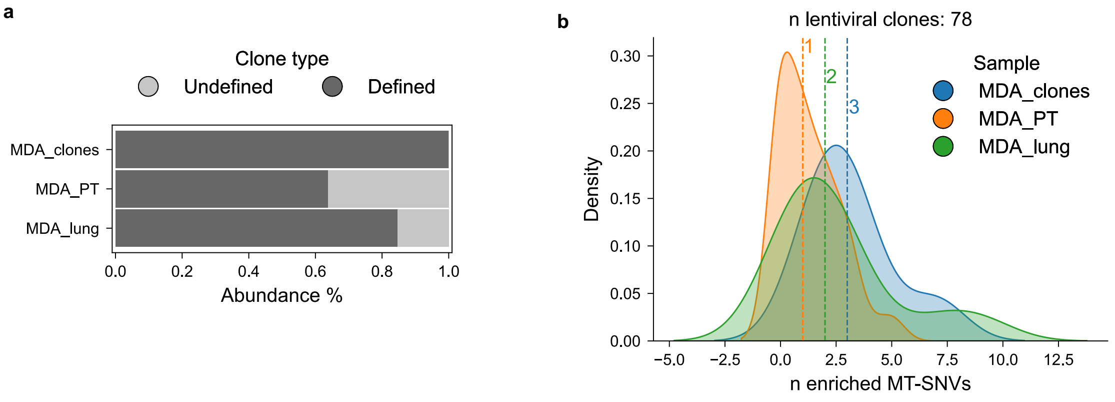


**Supp. Fig. 10. Supervised characterization of ground-truth clones MT-SNVs load. a.** Relative abundance of GBC-clones with at least one – or without any - clonally enriched MT-SNV - i.e., genetically defined and undefined clones. **b.** Density plot showing the sample-wise distribution of GBC-clones MT-SNVs load – i.e., the number of clonally enriched MT-SNVs, for each clone.

#### MT-genotyping

Accurate definition of binary (0: wild-type, 1: mutated) MT-genotypes is a fundamental step of MT-scLT workflows, as algorithms for clonal – e.g., `leiden` algorithm – and phylogeny inference – e.g., `UPMGA`, `NJ`, `mpboot`, `iqtree` – can take as input cell-cell distances – or kNN graphs, representing genetic dissimilarity with discrete metrics like jaccard or hamming distance – or cell profiles represented as discrete strings of characters – reference or alternative alleles. To benchmark our `MiTo` genotyping algorithm against the simpler `vanilla` approach – `--bin_method` hyper-parameter in `nf-MiTo`, see also Supplementary Information – `MiTo` toolkit – `MiTo` – MT-genotyping – we started from cell-filtered `maegatk` AFMs as shown before. First, tuned the internal parameters of `MiTo` genotyping (Supp. Fig. 11): i) the binomial mixture probability threshold – i.e., the decision threshold that assign the mutated/wild-type genotypes to a given cell and variant, given mutated and wild-type posterior probabilities –; and ii) the minimal prevalence required for a MT-SNV to be genotyped using binomial mixtures - `--t_prob` and `--min_cell_prevalence` hyper-parameters, respectively, see Supplementary Information – `MiTo` toolkit – `nf-MiTo` hyper-parameters. Different Probability thresholds – 0.5-0.09 - did not impact clonal inference accuracy, while probabilistic genotyping of MT-SNVs with less than 0.05 prevalence reduced accuracy, especially for the MDA_PT dataset. Therefore, we set default values to 0.7 - `t_prob` - and 0.05 - `min_cell_prevalence`.

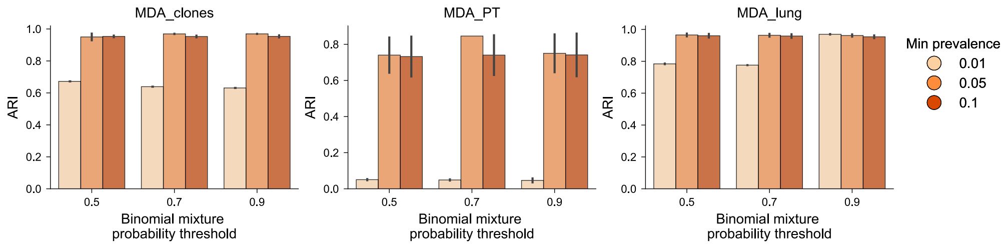


**Supp. Fig. 11. `MiTo` genotyping hyper-parameter tuning.** Adjusted Rand Index (ARI) for `MiTo` clonal inference after `MiTo` genotyping with variable values of binomial mixture probability threshold and minimum prevalence of variants genotyped with binomial mixtures, across samples.

Then, we benchmarked `MiTo` and `vanilla` (Supp. Fig. 12), requiring either more than 1 or 2 AD counts to assign the "mutated" genotype – `--min_AD` hyper-parameter in `nf-MiTo`, see also Supplementary Information – `MiTo` toolkit – `MiTo` – MT-genotyping. – Compared to `vanilla` - `MiTo` consistently achieved superior clonal inference accuracy across all samples, as measured by `ARI`, `NMI` and `AUPRC` metrics - with MDA_lung `AUPRC` as the only exception to this trend. Notably, within samples, requiring at least 2 consensus UMIs for the "mutated" genotype resulted in better accuracy, with negligible effect for MDA_lung but significant improvement in MDA_clones and MDA_PT datasets. The same trend was observed for the correlation between tree- and character-based distances. `MiTo` genotyping was more conservative in assigning the mutated state in the median number of MT-SNVs per cell in the MDA_clones sample, and by the slightly lower number of MT-SNVs retained in the final genotyped AFMs after the last filters implemented in `MiTo` MT-SNVs filter, after genotyping – see Supplementary Information – `MiTo` benchmark – MT-SNVs filtering. Consistently, `MiTo` genotyping resulted in slightly lower number of analyzed cells and GBC-clones, but without altering the overall clonal structure of the data. To further test the robustness to noise of the two genotyping strategies, we systematically perturbed observed AD counts. To this end, we perturbed each MT-site AD counts by adding spurious counts, sampled from binomial distributions with increased rate parameters. We simulated these perturbations varying either the binomial rate parameter (0.1-5x the fitted binomial rate parameter of the perturbed MT-site), or the number of MT-sites affected (10-100% filtered sites), performing 100 simulations for each perturbation condition. We quantified the effect of these perturbations considering cell-cell distances in the resulting MT-SNV spaces.

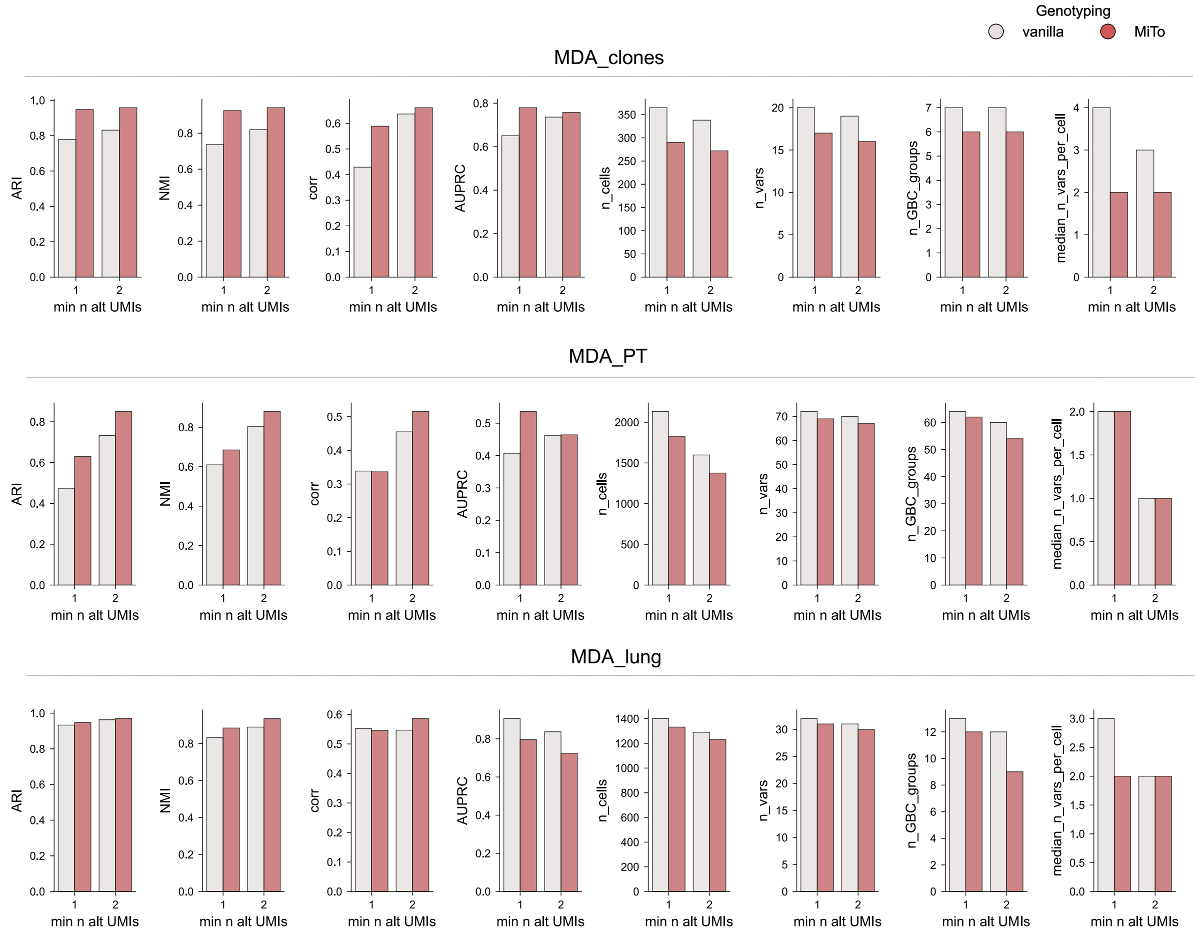


**Supp. Fig. 12. MT-genotyping benchmark.** `nf-MiTo` metrics – i.e., clonal inference accuracy (ARI, NMI, AUPRC), cellular yield (n_cells, n_vars, n_GBC_groups, median_n_vars_per_cell) and tree structure (corr, correlation between tree-based and character-based cell-cell distances – after MiTo or vanilla genotyping, varying the minimum numer of alternative UMIs required to assign the "mut" genotype. See Supplementary Information – `MiTo` toolkit – `MiTo` genotyping and Supplementary Information – `MiTo` toolkit – `nf-MiTo` metrics.

Specifically, we quantified the level to which GBC clonal labels could be discriminated from cell-cell distances in MT-SNVs space with `AUPRC`, and the signal-to-noise ratio of MT-phylogenies with the correlation between tree- and character-based distances - see Supplementary Information – `MiTo` toolkit – `nf-MiTo` metrics. `MiTo` consistently achieved higher `AUPRC` and tree- character-based distance correlation.

Together, these results demonstrated the practical utility of our probabilistic approach to precisely assign MT-genotypes, minimizing False Positive mutated genotypes, and highlight its utility in presence of noisy data, a common scenario in single-cell data analysis.

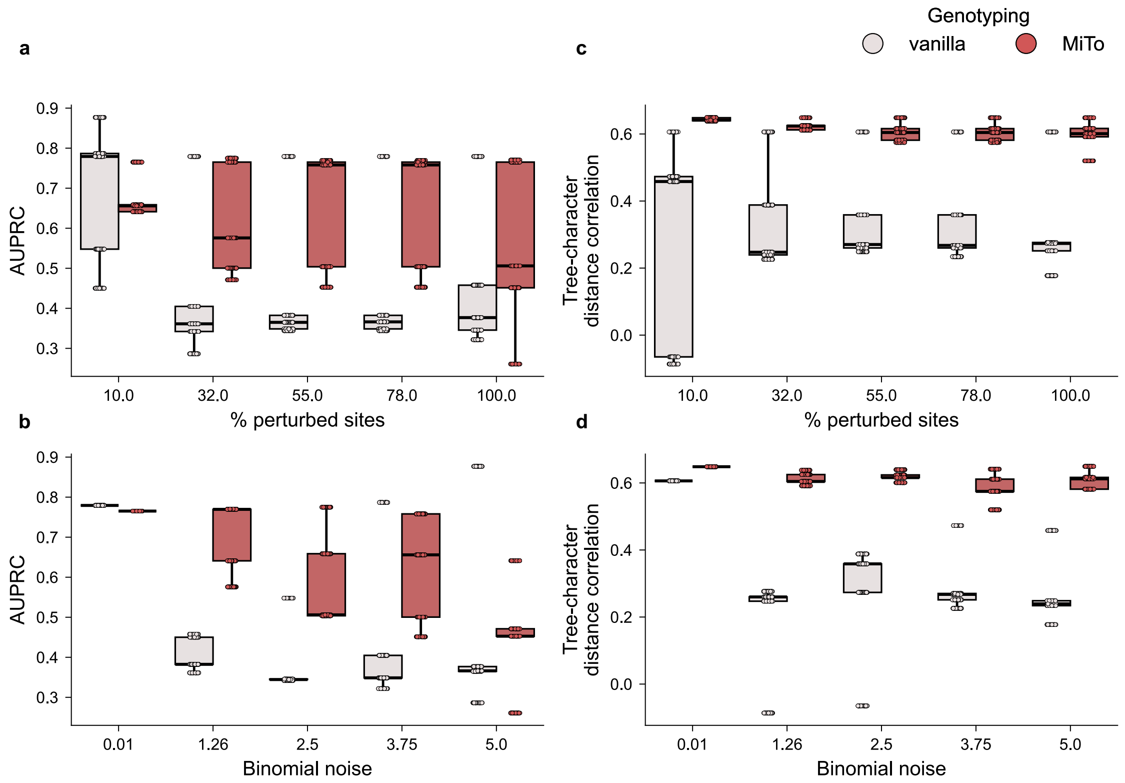


**Supp. Fig. 13. `MiTo` vs vanilla genotyping with simulated noise in alternative allele (AD) UMI counts. a-b.** Boxplots representing AUPRC values with variable % of perturbed sites (**a**) and level of binomial noise introduced in AD counts (**b**). **c-d** Same as in a-b, but for tree- character-based distances correlation – see Supplementary Information – `MiTo` toolkit – `nf-MiTo` metrics.

#### Distances in MT-SNV space

Several algorithms for clonal and phylogeny inference are based on pair-wise cell-cell distances in some genetic space. Thus, accurate quantification of these distances can directly impact lineage inference performance. `nf-MiTo` supports different metrics to quantify distances in MT-SNV space, with the jaccard distance being recently used successfully by [8]. Specifically, Weng et al. developed a modified a weighted version of this metric, with each MT-SNV weight inversely proportional to the mean prevalence of the variant across their patients' cohort. This idea was inspired by other strategies in scLT field, and can be regarded as a practical attempt to reduce the impact of common technological artifacts/non-somatic MT-SNVs. However, this approach is limited in its applicability by the need of multiple, independent samples, and implies that this kind of spurious characters have not been filtered out from the genotyped AFM – something that the `MiTo` MT-SNV filter is able to take care of, as demonstrated in Supplementary Information – `MiTo` benchmark – MT-SNVs filtering. For a more general approach, we tested an alternative weighting strategy. The unweighted jaccard distance is a strict measure of set between 2 binary MT-genotype profiles. As such, each positive MT-SNVs is treated equally, regardless of each variant detection strength. `MiTo` weighted jaccard distance weights each MT-SNV by its median allelic frequency in mutated cells – a value that represents the molecular evidence supporting these detection events. When we benchmarked standard – i.e., unweighted – jaccard vs weighted jaccard distances (Supp. Fig. 14), we scored very similar performances for nearly all metrics, with higher of ARI and NMI but lower AUPRC and tree- vs character-based correlations for the weighted distance compared to the unweighted one. Thus, better quantification of cell-cell distances in MT-SNV spaces will need further tuning in future works.

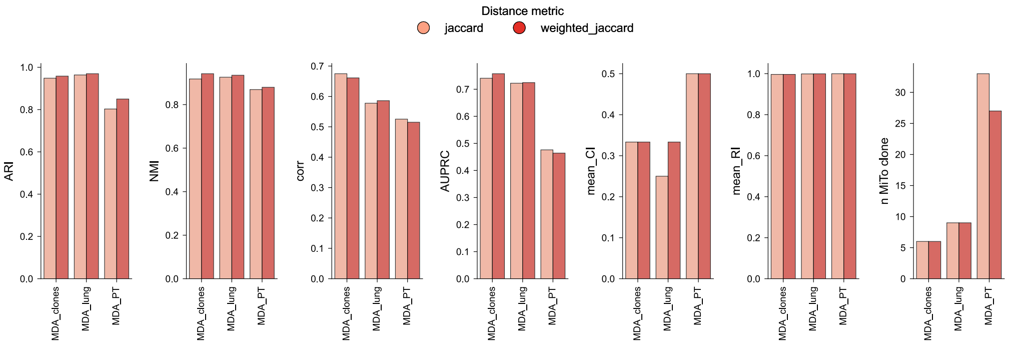


**Supp. Fig. 14. Distance metric benchmark**. Lineage inference performance - measured by selected metrics, see Supplementary Information – `MiTo` toolkit – `nf-MiTo` metrics – compared between jaccard and weighted jaccard distances.

#### Clonal inference

The final, fundamental step of `nf-MiTo` is lineage inference, in terms of both phylogeny and MT-clones. The `MiTo` approach to infer discrete clonal labels by efficiently "cutting" MT-phylogenies in disjoint, MT-SNVs-supported clades is described in Materials and Methods – Computational Methods – `MiTo` – Clonal inference from MT-phylogenies. Here we will provide additional information about the 3 clonal inference algorithms that we benchmarked `MiTo` against, and the benchmark itself.

<u>Other Methods</u>

`vireoSNP` is a Bayesian clustering algorithm designed for demultiplexing pooled single-cell RNA-seq (scRNA-seq). Its input consists of allelic read or UMI counts at selected SNPs across single cells - typically obtained via cellSNP-lite - and optionally the genotypes of donors.

vireoSNP models observed counts with a variational binomial mixture model, where donor genotype and cell assignment probabilities are jointly estimated by posterior maximization. vireoSNP allows flexible prior specification, including known donor number (K) and genotypes – selected by visual inspection of the Evidence Lower Bound (ELBO) across multiple K values. Typical outputs include donor assignment per cell, inferred genotypes, and assignment confidence scores. Originally developed for sample multiplexing, doublet detection, and quality control in large scRNA-seq studies, it has also been applied to clonal inference from MT-data.

The leiden algorithm is a graph-based clustering method widely used in a variety of single-cell omic settings to identify discrete cell clusters. It operates on k-nearest neighbor (kNN) graphs, where nodes represent cells and edges reflect similarity. The method partitions the graph by optimizing modularity, improving upon its predecessor – i.e., the louvain algorithm by ensuring well-connected communities through iterative refinement. The key hyper-parameter parameter here is the resolution parameter, which controls cluster granularity. Widely adopted in popular GEX single-cell workflows - e.g., Scanpy or Seurat -, leiden is usually applied after data normalization, dimensionality reduction, and neighborhood graph construction. Its output is a discrete cluster label for each cell. Its efficiency and ability to capture biologically meaningful clusters make it a gold standard for unsupervised clustering in scRNA-seq analyses. Together with the louvain algorithm, it is arguably the most popular algorithm for clonal inference from MT-data.

CClone is a Nonnegative Matrix Factorization (NMF) method developed for clonal inference from single-cell mutational data – nuclear and MT- SNVs. The input consists of a filtered variant call matrix M and an associated weight matrix W that captures per-cell, per-variant coverage confidence. CClone applies a weighted NMF matrix decomposition where observed M entries are weighted and missing entries - due to lack of coverage - are ignored. To select the optimal number of clones (K hyper-parameter), CClone evaluates a range of K values and selects the result with the highest orthogonality score among cell factors, indicating anti-correlated cell factors. These cell factors can be used for downstream clonal assignment, while variant factors can be used to detect clone-specific SNVs.

Of note, we have not included in this benchmark 2 other clonal inference methods specifically developed for MT-data: LINEAGE[9] and CloneTracer[10]. Both methods were developed to handle lower number of cells and variants that the ones presented in this work, and were not able to complete inferences in reasonable time or went out-of-memory – on a 28 CPUs work-station with 189 GB of RAM. For these reasons we were unable to evaluate their performance, and we excluded from the benchmark.

MT-SNV space selection

To test the performance of each clonal inference method on MT-SNV spaces – i.e., some filtered and genotyped AFM – with reasonable phylogenetic signal, we proceeded as follows. First, for each dataset we filtered the hyper-parameter combinations that yielded:

- >250 cells and >6 GBC clones for the MDA_clones dataset
- >1000 cells and >30 GBC clones for the MDA_PT dataset
- >1000 cells and >10 GBC clones for the MDA_lung dataset

and fraction of unassigned cells <10%. Then, for each sample we selected only the top hyper-parameter combinations (n=5) ranked by ARI – evaluating concordance between GBC- and MT-clones inferred with the `MiTo` clonal inference approach. These MT-SNV spaces were used to test all the other methods.

Tuning methods hyper-parameter

Each method was optimized in terms of key hyper-parameters. For `vireoSNP`, we used the same procedure described here, but making the selection of the optimal K value automatic. Specifically, for each K we computed ELBO values for n=50 independent runs of the algorithm, and selected the K value at the knee of the curve described by the median – across runs - ELBO value (y-axis) for each K value (x-axis). Selected the optimal K, a final clonal inference run was performed to yield final cell assignment probabilities. Each cell was assigned to the clone for which the assignment probability was higher - >0.7, and labelled as "unassigned" otherwise. The K parameter ranges tested were: 2-15 for MDA_clones, 2-60 for MDA_PT, and 2-20 for MDA_lung.

For the `leiden` algorithm, we first derived a kNN graph from each complete cell-cell distance matrix (k=15). This graph was partitioned with n=50 resolution values - range 0.5-2.5. The mean silhouette score vas used to select the best clustering resolution – i.e., the resolution value providing clonal labels with the highest mean silhouette scores evaluated on the full pre-computed cell-cell similarity matrix – weighted jaccard distances between cell MT-genotypes. For `CClone,` K was varied as done for `vireoSNP`, and for each run, the orthogonality score between estimated wNMF cell factors was coumputed. The K value with highest orthogonality score vas selected for each dataset, and each cell was assigned to the factor – i.e., MT-clone - for which `CClone` scored the highest value.

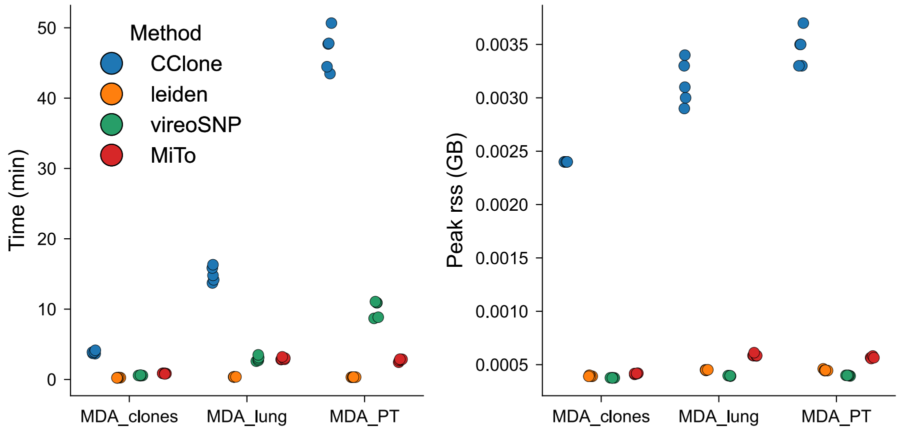


**Supp. Fig. 15.** Time (left) and peak memory (right) requirements for the clonal inference benchmark – grouped by dataset and method.

### Multi-omic analysis of longitudinal Breast Cancer clones

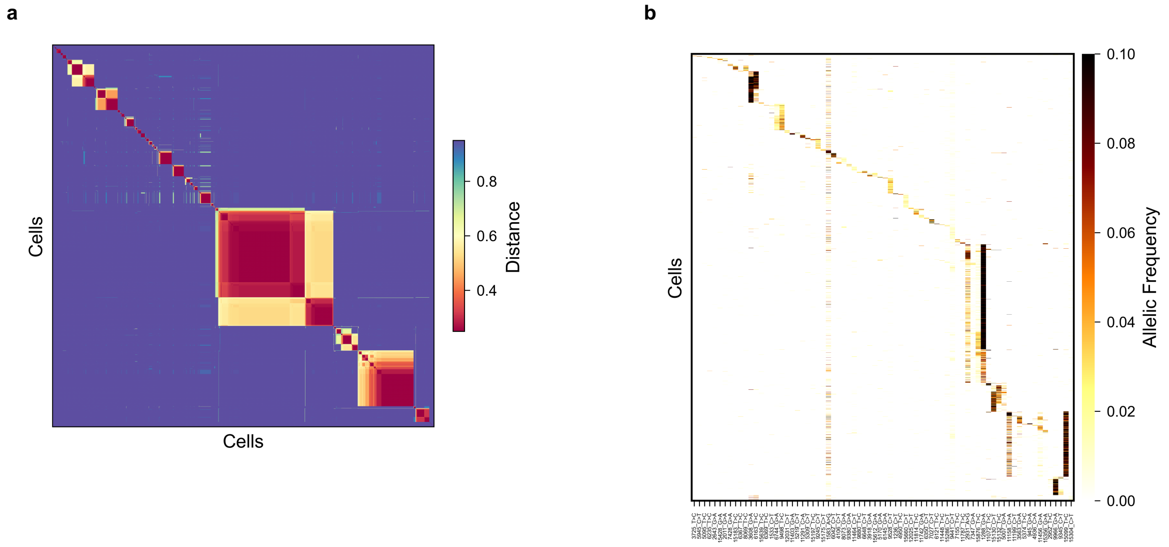


**Supp. Fig. 16. Lineage inference in longitudinal PT-lung dataset. a.** Clustered cell-cell distances (weighted jaccard distance) in MT-SNV space. Cells (n=2549) are ordered as in the UPMGA phylogeny in **Fig. 4c**. **b.** Cell (2549) x variant (77) AF heatmap. Cells (rows) ordered as in **b**, MT-SNVs are ordered according to their clone enrichment value – see Materials and Methods – Computational Methods – `MiTo` – Clonal inference.

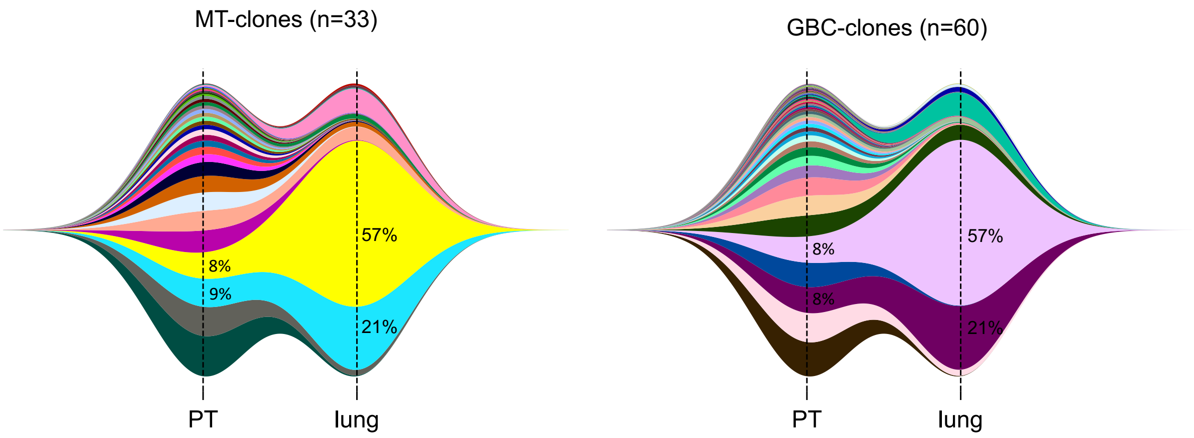


**Supp. Fig. 17. MT- and GBC- clonal prevalences in the PT-lung dataset.** Fishplots representing clonal prevalences in PT and lung samples for MT- (n=33) and GBC- (n=60) clones. Selected clones are annotated across timepoints.

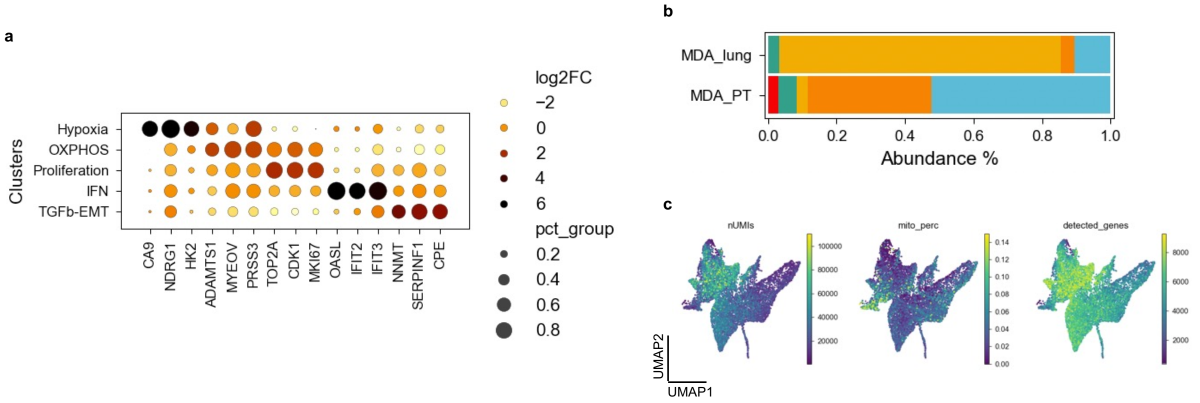


**Supp. Fig. 18. Cell state annotation and abundance in PT-lung dataset. a.** Dotplot with selected marker gene expression (columns) across annotated cell clusters (i.e., cell states in Fig. 4b-c, rows). **b.** Cell state abundance across PT and lung datasets – color-code as in Fig. 4b. c. Technical covariates plotted unto UMAP cell embeddings – same as in Fig. 4b.
